# Knowledge Graphs and Explainable AI for Drug Repurposing on Rare Diseases

**DOI:** 10.1101/2024.10.17.618804

**Authors:** P. Perdomo-Quinteiro, K. Wolstencroft, M. Roos, N. Queralt-Rosinach

## Abstract

Artificial Intelligence (AI)-based drug repurposing is an emerging strategy to identify drug candidates to treat rare diseases. However, cutting-edge algorithms based on Deep Learning (DL) typically don’t provide a human understandable explanation supporting their predictions. This is a problem because it hampers the biologists’ ability to decide which predictions are the most plausible drug candidates to test in costly lab experiments. In this study, we propose *rd-explainer* a novel AI drug repurposing method for rare diseases which obtains possible drug candidates together with human understandable explanations. The method is based on Graph Neural Network (GNN) technology and explanations were generated as semantic graphs using state-of-the-art eXplainable AI (XAI). The model learns features from current background knowledge on the target rare disease structured as a Knowledge Graph (KG), which integrates curated facts and their evidence on different biomedical entities such as symptoms, drugs, genes and ortholog genes. Our experiments demonstrate that our method has excellent performance that is superior to state-of-the-art models. We investigated the application of XAI on drug repurposing for rare diseases and we prove our method is capable of discovering plausible drug candidates based on testable explanations. The data and code are publicly available at https://github.com/PPerdomoQ/rare-disease-explainer.

**Highlights:**

- We demonstrated the use of graph-based explainable AI for drug repurposing on rare diseases to accelerate sound discovery of new therapies for this underrepresented group.
- We developed *rd-explainer* for rare disease specific drug research for faster translation. It predicts drugs to treat symptoms/phenotypes, it is highly performant and novel candidates are plausible according to evidence in the scientific literature and clinical trials. Key is that it learns a GNN model that is trained on a knowledge graph built specifically for a rare disease. We provide *rd-explainer* code freely available for the community.
- *rd-explainer* is researcher-centric interpretable ML for hypothesis generation and lab-in-the-loop drug research. Explanations of predictions are semantic graphs in line with human reasoning.
- We detected an effect of knowledge graph topology on explainability. This highlights the importance of knowledge representation for the drug repurposing task.

## 1. Introduction

Developing new drugs can be a challenging effort that often ends with the drug not being able to launch. Recent studies have shown that around 90% of drugs fail to be approved during their clinical development [1]. This leads to a fruitless expenditure of both time and money that will yield no financial returns. The situation is even worse in the case of rare diseases, as pharmaceutical companies may consider it risky to invest large amounts of resources into developing drugs that only a small percent of the population will need. Nonetheless, in total, human beings are affected by approximately 7,000 rare diseases, of which only 5% have an effective treatment [2]; and only in Europe about 36 million people suffer from rare diseases [3].

In this scenario, drug repurposing strategies have appeared as a possible approach to solve these issues. By reusing drugs that have already been approved, companies can avoid many of the costly and time-consuming steps of clinical trials. In this context, innovative approaches to drug repurposing, such as computational strategies and AI-driven methodologies, have emerged as promising solutions to address these challenges. Graph-based drug repurposing is another noticeable strategy that has gained attention in recent years. By constructing intricate networks of molecular interactions, genes, proteins, and diseases, this approach unveils hidden relationships and connections that might otherwise go unnoticed [4].

Still, many people remain skeptical about AI-driven decisions, specially Machine Learning (ML) and Deep Learning, as many of them come with no explanation that can help to understand the reason why they should be trusted (also called black-box AI). This issue is especially significant in the healthcare field, where decisions may have an important impact on people’s lives. Also, giving valid explanations can help researchers to point in the right direction in the generation of hypotheses that are testable in the lab and enable a solid knowledge discovery. Furthermore, the EU General Data Protection Regulation (GDPR) is requesting the AI industry to fulfill the ’right to explanation’ [5]. This ’right to explanation’ implies that when a decision is significantly affected by an automated process/algorithm, the individual can demand an explanation. In recent years, many different tools have appeared to try and cover this gap in the emerging explainable AI (XAI) research area [6, 7, 8].

In this study, we explore whether AI can be used to produce both predictions and explanations in computational drug repurposing for rare diseases and, if so, how helpful can these explanations be for hypothesis generation. The main objective of this work was to develop and implement a pipeline to find marketed drugs that can be used to treat the symptoms of a rare disease. Our approach is based on cutting-edge AI algorithms used in computational drug repurposing such as graph ML using knowledge graphs (KG) and graph neural networks (GNN), and XAI methodology to provide the explanations supporting the drug predictions made by the AI model. The approach was evaluated by selecting Duchenne muscular dystrophy (DMD) as a case study, a genetic disorder that is the most common form of muscular dystrophy [9]. We demonstrate the generalizability of our approach by applying the pipeline to different rare diseases.

## 2. Related work

### 2.1. Knowledge graph-based drug repurposing

The state-of-the-art of computational drug repurposing approaches make use of graph-based structures and AI techniques to find potential drug candidates. One of the main advantages of using graph structures is that they can easily incorporate information from different sources. This is especially important in the domain of rare diseases, where information is distributed and often scarce. The ability to integrate as much relevant data as possible can confer a significant advantage. An example of this would be the recent study of Al Saleem et al. [10], where a knowledge graph was used to discover drug candidates to treat COVID-19.

Different ML algorithms can be used to analyse knowledge graphs, including matrix factorization, random-walk approaches (node2vec [11]), geometric embeddings (DistMul [12]) and GNNs [13, 14], each one of them with its own advantages and disadvantages, see Table 1. In our study, we used a combination of random-walk approaches and GNNs as in contrast to other methods (like matrix factorization or geometric embeddings) they can easily incorporate new information without the need of retraining the ML model. This is especially relevant in the field of drug repurposing where new information about drugs, genes and diseases is being published [15, 16, 17].

**Table 1.** Table showing the main advantages and disadvantages of different graph ML methods.

| Method | Example | Advantages | Disadvantages |
| --- | --- | --- | --- |
| *Matrix factorization* | ADA-GRMFC [18] | Simple Model<br>Global Information | Computationally expensive<br>Scalability<br>Difficult to include new values |
| *Random-walks* | node2vec [11] | Simple model<br>Computationally efficient | Only local information<br>Can't learn from features<br>Can't capture structural information |
| *Geometric Embeddings* | DistMult [19] | Interpretability<br>Good performance | Only local information<br>Can't capture structural information |
| *GNNs* | GraphSAGE [20] | Global and local information<br>Structural information | Black-box<br>Computationally expensive |

### 2.2. Explainable AI on graph ML

One of the graph-based methods that can provide explanations of the predictions, also called local explanations, is (Graph)LIME [6], an adaptation of the popular and more general explainability method LIME [7]. The idea behind this method is the following: when trying to get an explanation for a given prediction, (Graph)LIME performs small perturbations to the features of nodes, and sees how the predictions vary with respect to the initial prediction. The more the prediction changes, the more the model is relying on that feature to obtain its prediction. This way, explanations in this model are given in the form of a set of node features. Among its drawbacks, this method can only be used in node classification tasks. Another explainability method is CRIAGE [8] where explanations are given as a set of rules.

Finally, the method chosen in this work is GNNExplainer [21]. The insight of how this method works is the following: given an initial prediction (link prediction, node classification or graph classification) obtained through a GNN, GNNExplainer finds a subset of node features and edges that are responsible for the prediction. This subset is obtained by training an edge and node mask. This method was chosen as explanations are provided in the form of a subgraph that can be easily understandable. Additionally, it is model-agnostic, which means that if more sophisticated GNNs are developed in the future, these new GNNs can be easily incorporated into the pipeline. These features make it a popular method in the research community [22, 23, 24]. However, a major drawback is that it lacks consistency when obtaining explanations. This means that explanations on the same prediction can significantly change if running GNNExplainer several times.

## 3. Methods

### 3.1. *rd-explainer* method overview

*rd-explainer* is the drug repurposing method we developed for rare diseases and its pipeline is illustrated in Figure 1. rd-explainer has three modules: the Knowledge Graph Construction module constructs a KG for the specific rare disease and drug repurposing task, the Prediction module trains a GNN model and predicts drug candidates for the rare disease symptoms, and the Explainer module computes the most important semantic subgraphs that explain the connection between the predicted drug and the symptom. Firstly, information related to the disease is gathered from different data sources: Monarch Initiative knowledge base [25] for disease pathology, and DrugCentral [26] and Therapeutic Target Database [27] for disease druggability. This information is then preprocessed and captured as a knowledge graph. Next, for each node in the graph a feature vector is obtained that will be used as input for the GNN model. This is done by making use of a method known as edge2vec [28] to consider the different edge semantics in the KG for node embedding learning. We used the version extracted from Github (accessed in 2021) ^1^. The following step is to build and train the GNN model, which is done using the GraphSAGE framework for graph representation learning [20]. Next, it is the link prediction for each drug-symptom node embeddings pair by using the dot product as scoring function. Finally, we produced prediction explanations as semantic graphs using GNNExplainer [21], a recent and, to our knowledge, one of the first XAI methods for obtaining explanations from GNN predictions. GraphSAGE and GNNExplainer were implemented using using Pytorch Geometric version 2.0.4.

**Figure 1:**
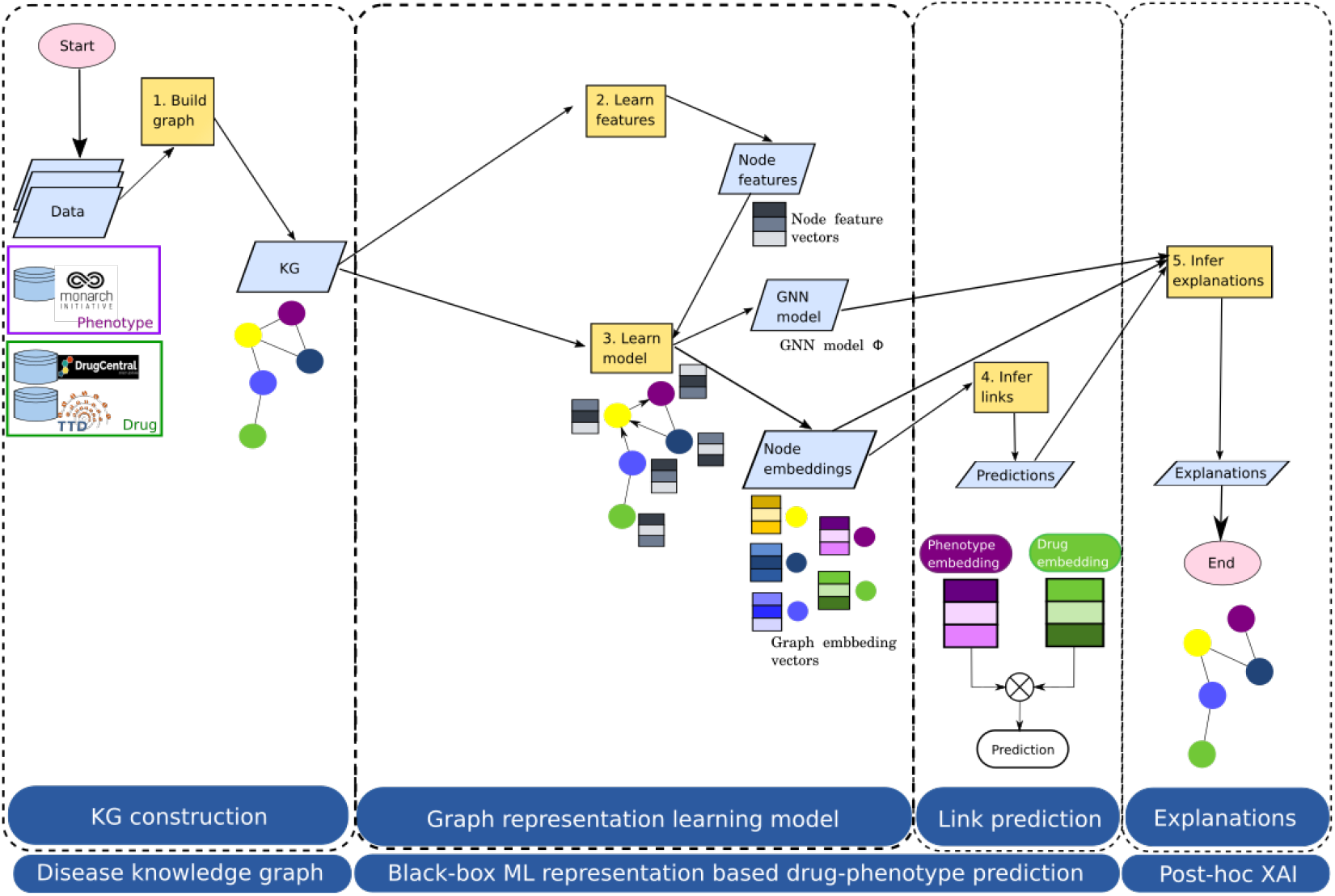
*rd-explainer* drug repurposing method pipeline developed in this work.

The code is freely accessible with an open license at https://github.com/PPerdomoQ/rare-disease-explainer.

### 3.2. Rare disease-specific drug repurposing knowledge graphs

#### 3.2.1. Data sources

Data was obtained from three different sources: Monarch s[25] (accessed in 2021), DrugCentral [26] (2021 version) and Therapeutic Target Database (TTD) [27] (November 8th, 2021 version). Monarch is a knowledge base built on semantic principles, unifying gene, variant, genotype, phenotype, and disease data across different species. Its primary aim is to establish links between genes and phenotypes, thereby facilitating computational exploration of human disease biology. Monarch was chosen as it contains curated information across different species. This way, because rare diseases are often less studied than common diseases, incorporating information from other species can maximize the amount of knowledge in the graph. However, Monarch does not specialize in drug information.

Drug information was incorporated from DrugCentral (drug-target information) and from Therapeutic Target Database (drug-disease information). DrugCentral is a comprehensive online database that provides information about approved drugs, active ingredients and other pharmaceutical products. One of its major features is that it is open source and its data is freely available to anyone. For this project, we made use only of the drug-target information (as it is the main piece of information that is not present in Monarch) downloaded as a tsv file from their site [27] ^2^. Similarly, TTD is a database that specializes in drugs and their respective therapeutic targets. Once more, this database is freely accessible and its information can be easily downloaded in csv format (in this project, we just made use of the drug-disease information [27] ^3^, once again because it is the information that is missing in Monarch).

#### 3.2.2. Knowledge graph construction

To extract information from Monarch, the BioKnowledge Reviewer [29] tool version 1.0 was used. This tool was originally created to collect knowledge from several sources and create a knowledge graph that could be later used for hypothesis generation. It works by using several seeds (node identifiers (IDs)) as input to query the Monarch API and constructing the graph based on the neighborhood of those seeds. After introducing the seeds in the BioKnowledge Reviewer pipeline, the final output is the rare disease research question specific knowledge graph structured in two dataframes (stored as csv files). One of them contains a list of nodes with their respective name, IDs, semantic entity type, synonyms and description. The second file contains the list of edges, again containing the IDs of the entities participating in each link and other edge information such as type of edge, supporting evidence and reference date. Monarch was our main source of information, and so it served as a starting point to create the rest of the graph. This way, data from other data sources was modified to fit Monarch’s standards. Finally, the graphs were constructed using the networkx Python library [30] version 2.3.6. With this library the dataframes extracted using BioKnowledge Reviewer were converted into a *Graph* object. For a more detailed understanding of the knowledge graph generation process, please consult the associated code available at: https://github.com/PPerdomoQ/rare-disease-explainer.

We integrated data in two different knowledge graphs to perform the experiments. Each one of them was constructed using different (number of) node seeds to extract information from Monarch. The first one (KG A) only uses two seeds: DMD seed (HGNC:2928), corresponding to the human gene that causes the disease; and DMD seed (MONDO:0010679), corresponding to the disease itself. The second graph (KG B), extends KG A by including as seeds all phenotypes of the rare disease (in total, 27 more seeds). The seeds used for the construction of each graph can be found in Tables S1 and S2. The idea of creating two different graphs is to find out if the performance of the model and the quality of the explanations increases by incorporating more (phenotypic) information.

### 3.3. ML model and XAI

#### 3.3.1. Node features

At this point, none of the nodes have any specific node features. It is possible to run a GNN relying only on graph information, i.e., network topology (this is done, for example, by using the node degree as graph feature); nonetheless, this resulted in a poor performance (results not shown). To increase the efficiency of the model, edge2vec was used to produce a specific embedding for each node that captures information about its neighborhood. edge2vec [28] is a tool that generates node embeddings based on the node neighborhood and types of edges connecting each node. After executing edge2vec, each node was given a unique feature vector.

#### 3.3.2. Data splitting

As any other machine learning task, data needs to be splitted into training, (validation) and test sets. However, when tackling a link prediction task, there are different ways to perform this split. In link or edge prediction tasks, edges can be divided into two groups: message passing edges and supervision edges. Message passing edges are the ones that will be used by our GNN to obtain the embeddings, while supervision edges are the ones that will be used to test the performance of our model [31, 32]. Additionally, when creating the supervision edges it is necessary to include negative examples by applying negative sampling. These negative sample edges are edges that are not present in our original graph, i.e., entities that it is known are not linked or there is no known link between them, and the idea is that the neural network is able to learn to distinguish true or positive edges from false or negative edges. In general, one negative edge is created for each true edge [31, 32].

In this work, the method that was selected was the all-graph transductive split [31, 32]. When applying this method the division is done in the following way: in the training dataset the supervision edges and the message passing edges are the same; in the validation dataset the message passing edges are the training edges (message and supervision) and the supervision edges are different from the training supervision edges; finally, the test set message passing edges are formed by the validation edges and supervision edges that are different from the training and validation supervision edges.

This method is one of the standard settings when performing link prediction tasks, as the whole graph can be seen in all dataset splits [31]. The proportion used were 80% of edges used for training set, 10% for validation set and 10% for test set. The training set will be used to train the model, the validation set to select the best hyperparameters, and the test set to obtain the global performance of the model.

#### 3.3.3. GNN model

We first utilized a GNN algorithm to learn vector representation embeddings for nodes in our knowledge graphs. Then, we applied these node embeddings for drug-phenotype link prediction. The GNN algorithm that we used in this work is called GraphSAGE [20]. GraphSAGE performs inductive graph representation learning by leveraging rich node attribute information. The main advantage that was brought by GraphSAGE is its scalability: instead of working with full batches (the whole graph is seen during the training) it works with mini-batches. Each mini-batch is a subset of computational graphs (a computational graph is the individual GNN that is built for each node) of N nodes. By applying this technique, the GNN can better manage larger graphs. The GraphSAGE model was created using the DeepSNAP library [32] version 0.2.1 to obtain the predictions. The hyperparameter optimization was performed using RayTune [33] version 1.12.1, as it is a model-agnostic library that allows to run multiple trials in parallel, reducing the training time. The list of hyperparameters that were needed to be tuned and the optimal values can be found in Table S5. In total, 30 models were created (each of them containing a random selection of parameters).

#### 3.3.4. Drug-phenotype link predictions

The GNN model generates embeddings for individual nodes within the graph as its final output. By applying the dot product between distinct node pairs and applying a sigmoid function, we obtain a value that shows the likelihood of a link existing between those nodes. Consequently, we obtain dot products between each drug and every phenotype in the graph, and rank them in descending order. The top-ranked dot products are considered the most promising targets. Links that were already present in the graph were removed from the ranking.

#### 3.3.5. Graph-based prediction explanations

We applied GNNExplainer to generate explanations for every drug-phenotype prediction. To do so, we adapted the pipeline code (from Pytorch geometric version 2.0.9) to generate explanations for the link prediction task, which was not implemented in authors’ version [21] (see pseudocode in Algorithm 1 in the Supplementary material). However, this XAI algorithm has a problem of robustness in the explanations it produces [34] and, additionally, it may yield disconnected graphs affecting to the interpretability of explanations by domain-users. To solve this issue, we developed the following procedure. First, we make the assumption that a complete explanation is one that connects the two targeted nodes. If drug A can treat phenotype B, there must be some common pathway that allows A to interact with B. This way, the procedure starts by running GNNExplainer for several iterations. In each iteration, networkx is used to check if, in the subgraph generated by GNNExplainer, a path exists between both nodes. If no path is found, it continues with the next iteration; if it does exist, it stops iterating and that subgraph is considered to be the final explanation. If no subgraph is found that satisfies the ’pathway’ condition, the last subgraph is returned as a possible explanation.

In total 7 phenotypes were selected to evaluate the explanations (Muscular Dystrophy (HP:0003560), Respiratory Insufficiency (HP:0002093), Arrhythmia (HP:0011675), Congestive Heart Failure (HP:0001635), Dilated Cardiomyopathy (HP:0001644), Cognitive Impairment (HP:0100543) and Progressive Muscle Weakness (HP:0003323)). These phenotypes were selected to cover all the main areas that are affected by the disease (muscular, respiratory, cardiac and intellectual symptoms). For each prediction obtained in these phenotypes (three drug predictions per phenotype), an explanation was obtained. This process was done for the predictions coming from KG A and for those coming from KG B. This makes a total of 42 explanations (21 for each graph).

Regarding the parameters of GNNExplainer, because the graphs are highly connected, explanations were generated by using the 1-hop neighborhood around the graph. Using a higher k-hop neighborhood is not recommended as the amount of nodes in the subgraph increases exponentially which can make it difficult to understand the explanation. This happens because both graphs are scale-free graphs, and thus, by increasing the number of hops there is a higher chance that a ’hub-node’ is hitted, and the number of nodes escalates exponentially (see Section 4.1 in the results).

Additionally, the maximum size of the explanations was set to 15 (this means that no more than 15 edges will be part of the explanation). This way, we will avoid obtaining too complex explanations with many edges that might be impossible to comprehend by researchers. This was done by selecting the edges whose contribution values are among the 15th highest values.

Finally, the maximum number of iterations was set to 10. In other words, if after 10 iterations GNNExplainer has not found an explanations that connects the drug candidate with the targeted phenotype it will conclude that no ‘complete’ explanation was found, and the last explanation produced by GNNExplainer will be the one that will serve as final answer. This parameter can be increased or reduced depending on the expectations of the researcher. A large number of iterations increases the chances of finding a complete explanation at the cost of more computational time. On the contrary, reducing the number of iterations reduces the computational time, which can be useful if a researcher wants to obtain explanations for a large number of predictions.

### 3.4. Evaluation and metrics

#### 3.4.1. Evaluation of GNN model

##### Data

We used both graphs KG A and B. Data was splitted into three sets: training set, validation set and test set. **Baselines**. Our baselines include edge2vec [28], Graph-SAGE [20], ComplEX [35], DistMult [19], and TransE [36]. **Evaluation metrics**. The Area Under the Precision-Recall curve (AUPRC) was used to validate and test the performance of the model, as it has been shown that it leads to better precision when evaluating link prediction [37]. Additionally, we also computed the Area Under the ROC curve (AUROC), Precision, Recall and the F1-Score metrics - the harmonic mean of precision and recall.

Other evaluations were developed to further assess the performance of the model. These evaluations include the testing of different negative sampling sizes (n = 1, 5, 10 and 20) to determine the importance of keeping the data balanced. Additionally, both a regular 10-fold cross validation and a biased 7-fold cross validation were performed. The biased cross validation consists of the following: in each fold 4 phenotypes were removed from the training set, and it was observed how well the model was able to predict the links of the removed phenotypes.

#### 3.4.2. Evaluation of explanations

The evaluation of explanations was done manually, following a two-step process. Firstly, they were classified as complete or incomplete explanations based on the appearance of connection between the drug and the phenotype. We developed a function to visualize the explanations as semantic graphs (see section A.5 in the supplementary material for further details). This way, if the explanation contains a link between the drug and the phenotype it is considered to be a complete explanation. These explanations are considered the ones that are truly useful as they are the ones that can be easily understood and interpreted. On the other hand, explanations where there is no link between drug and phenotype (where there are two separate clusters) or where only one of the target elements (either the drug or the phenotype) is missing, are considered incomplete explanations. Several illustrative examples are provided in the supplementary material (See section A.6).

During the second step, we evaluated explanations using an objective and a subjective approach. First, complete explanations were reviewed and a manual search was performed to check whether the explanation proposed by the model had been already described in the literature (objective evaluation). This process was only performed for those predictions that have supporting evidence in the literature and that were classified as complete explanations. The examination of the literature was performed using PubMed and Google Scholar during the first half of 2022. Finally, each explanation was evaluated with our own biological knowledge (subjective evaluation).

## 4. Results

### 4.1. Rare disease KG topology and representation for drug repurposing

We generated two different drug repurposing knowledge graphs for the Duchenne muscular dystrophy rare disease. KG A contains 10786 nodes, 93905 directed edges. The average node degree of the graph 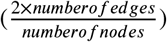 is 10.83, being the node with the highest degree, the human DMD gene, with a total degree of 1683. The diameter of the graph was 6, meaning that the longest shortest path between two nodes is 6 (in other words, one can travel from one node to another in 6 steps or fewer). The final feature that was obtained is the clustering coefficient, which measures the extent to which a graph is clustered together. In a complete graph (where all nodes are connected to all nodes) this clustering coefficient is equal to 1, while in a tree-like graph this coefficient is equal to 0. In KG A this clustering coefficient is equal to 0.33. A summary of the features can be found in Table 2.

**Table 2.** Table showing features of KG A and B.

| Property | KG A | KG B |
| --- | --- | --- |
| Number of Nodes | 10786 | 83665 |
| Number of Directed Edges | 93855 | 1984774 |
| Number of Undirected Edges | 58435 | 1440418 |
| Average Degree | 10.83 | 34.43 |
| Highest Degree | 1683 | 4817 |
| Diameter | 6 | 7 |
| Average Clustering Coefficient | 0.33 | 0.48 |
| Number of drugs | 337 | 1565 |
| Number of diseases | 5419 | 25636 |
| Number of drug-disease pairs | 86 | 599 |

In the case of KG B (built from 29 nodes: KG A seeds extended by 27 phenotypes of DMD), the total number of nodes is 83665, with a total of 1984774 directed edges. The average degree in this case is of 34.43, being the node with the highest degree the physiological process ’Protein Binding’ with a total degree of 4817. The diameter of the graph is of 7, which shows one of the features of scale-free networks: despite increasing the number of nodes 8 times and the number of edges 20 times, the diameter of graph B only increased one unit with respect to graph A. In this case, the clustering coefficient is equal to 0.48, showing that KG B is more clustered. Table 2 shows a summary of the features of both graphs.

The schema of the knowledge graph, which is the same for KG A and KG B, can be seen in Figure 2 and shows how the 8 different node types interact with each other. The schema contains 24 and 29 different edge types for KG A and KG B respectively, which are not included in this figure for clarity, but are listed in the Supplementary material S3 and S4.

**Figure 2:**
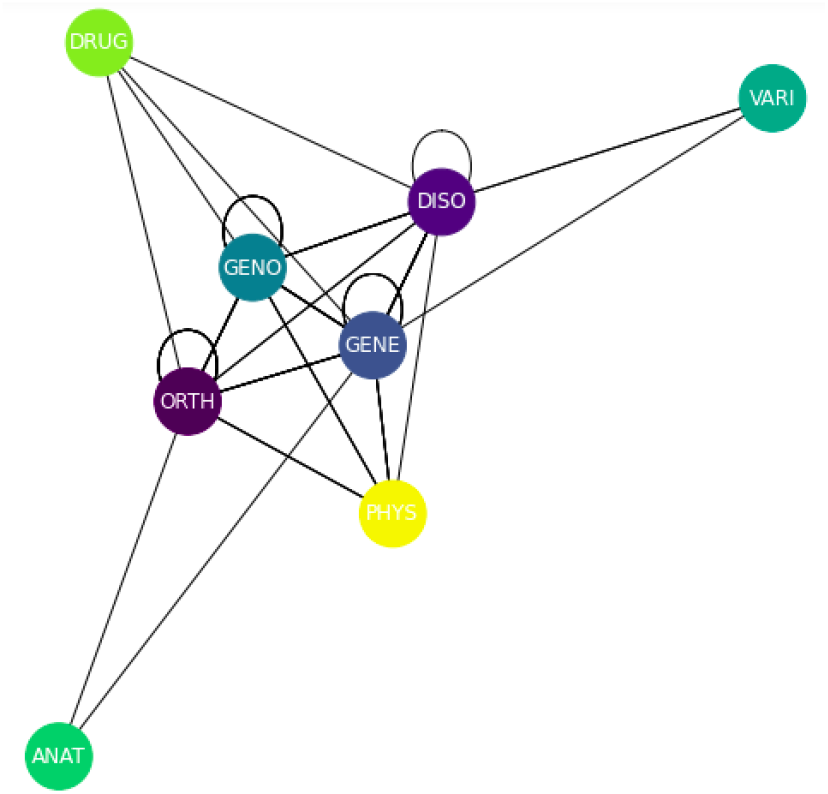
Schema of the knowledge graph. Node types are: drugs or chemical compounds (DRUG), genes (GENE), symptoms/phenotypes or diseases (DISO), gene variants (VARI), genotypes (GENO), gene orthologs (ORTHO), anatomical structures (ANAT), and biological processes (PHYS).

### 4.2. GNN model performance for rare disease specific drug repurposing

In total, two GNNs were used, one trained on KG A and one trained on KG B. The hyperparameter optimization was developed using RayTune and the optimal values can be found in Table S5. These hyperparameters were obtained by training several GNN models (Random Search) on graph A; and were later used to train a GNN model on graph B.

To measure link prediction performance, the scores obtained were precision, recall and the F1-Score, and can be found in Table 3 (the threshold used was 0.8). We found that both models (the one trained with KG A and the one trained with KG B) yield to high performance (F1-Score = 0.93 and 0.94 in KG A and B, respectively). To visualise the performance of the link prediction task, the ROC curve of KG A and KG B obtained on the test set can be found in Figure 3 and Figure 4, repectively.

**Table 3.** Precision, Recall and F1-Score obtained on each dataset, trained on each graph.

| Dataset | Precision |  | Recall |  | F1-Score |  |
| --- | --- | --- | --- | --- | --- | --- |
|  | Graph A | Graph B | Graph A | Graph B | Graph A | Graph B |
| Training | 0.93 | 0.96 | 0.96 | 0.93 | 0.95 | 0.95 |
| Validation | 0.93 | 0.96 | 0.93 | 0.92 | 0.93 | 0.94 |
| Test | 0.93 | 0.96 | 0.93 | 0.92 | 0.93 | 0.94 |

**Figure 3:**
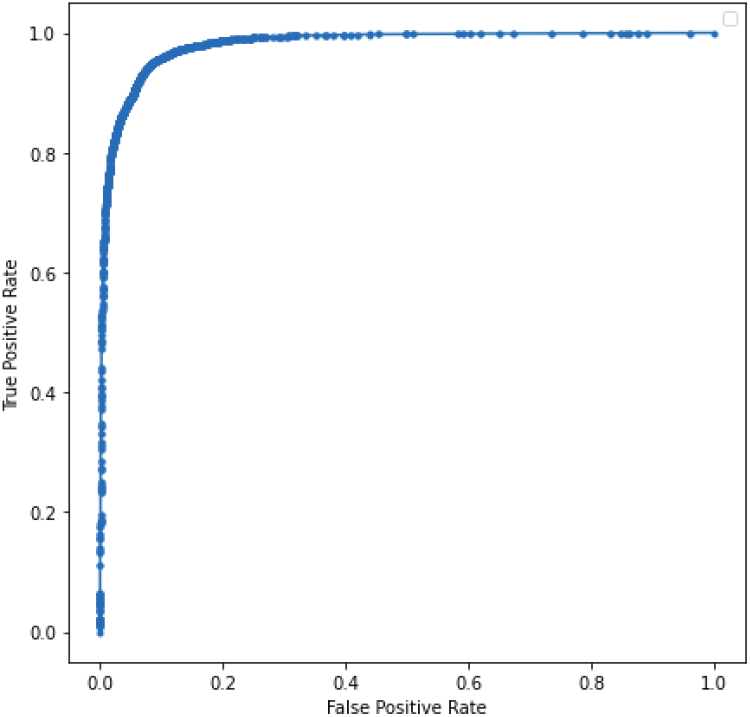
AUROC on the test dataset using KG A.

**Figure 4:**
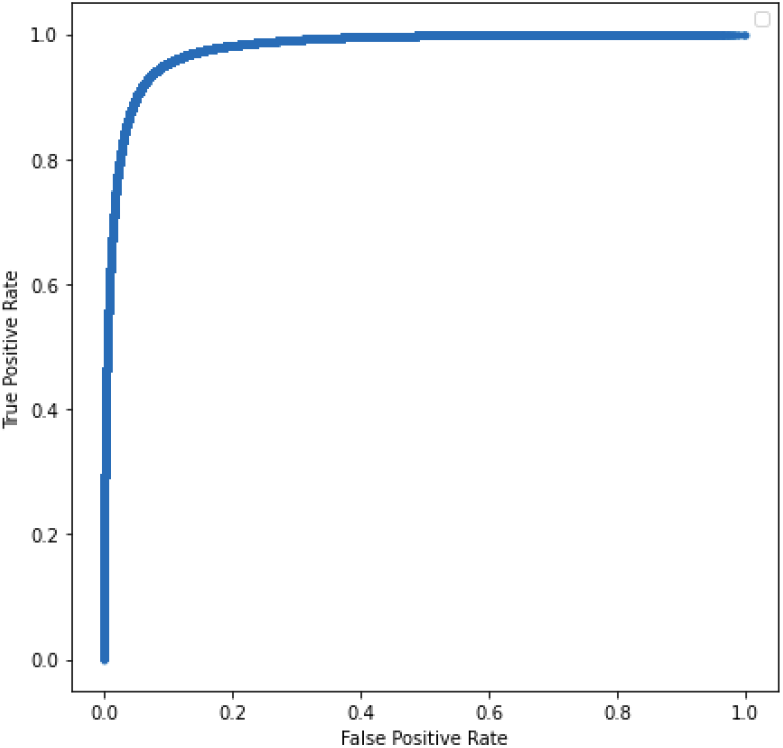
AUROC on the test dataset using KG B.

### 4.3. Evaluating rd-explainer with state-of-the-art methods

Firstly, we evaluated our GNN model applying different strategies and compared its performance to the state-of-the-art graph embeddings used in drug repurposing methods. Then, we evaluated our approach based on its ability to predict drugs that are already reported in the literature for a new symptom or phenotype.

We performed a regular 10-fold cross-validation and a biased 7-fold cross-validation evaluation in KG A. The regular 10-fold cross-validation obtained an average AUPRC of 0.98 and an average AUROC of 0.98. For the biased 7-fold cross-validation, in each fold 4 symptoms (along with the edges connected to those symptoms) were removed from the training set. Then the performance of the model was tested on the removed symptoms. In this case, the average AUPRC was 0.75 and the AUROC was 0.8.

The performance of the pipeline was evaluated for a different number of negative edges. This evaluation was only performed in KG A due to the large increase in the number of edges in the evaluation tests (and the consequential increase in the computational time). The results can be seen in Table 4. It is seen that as the number of negative edges increases, the PR curve is affected while the ROC curve remains mostly intact, a result that has previously been reported [38].

**Table 4.** Performance as the number of negative edge samples increases. This results were obtained using KG A.

| *Number of negative edges* | **Precision** | **Recall** | **F1-Score** | **AUROC** | **AUPRC** |
| --- | --- | --- | --- | --- | --- |
| 1 | 0.95 | 0.95 | 0.95 | 0.99 | 0.99 |
| 5 | 0.94 | 0.90 | 0.92 | 0.94 | 0.94 |
| 10 | 0.93 | 0.91 | 0.86 | 0.86 | 0.85 |
| 20 | 0.93 | 0.57 | 0.62 | 0.82 | 0.70 |

Finally, the performance of rd-explainer (tested in KG A) was also compared to other state-of-the-art methods, including edge2vec, GraphSAGE, ComplEX, DistMult, and TransE. Our results can be seen in Table 5 and they revealed that rd-explainer outperformed all the other methods based on the different evaluation metrics measured.

**Table 5.** Prediction performance metrics comparing rd-explainer with other state-of-the-art graph embedding methods including edge2vec, GraphSAGE, CompleEX, DistMult and TransE. The best results are **highlighted**. In the headings, **P** stands for *Precision*, **R** for *Recall*, and **F1** for *F1-Score*.

| *Method* | **P** | **R** | **F1** | **AUROC** | **AUPRC** |
| --- | --- | --- | --- | --- | --- |
| edge2vec | 0.90 | 0.90 | 0.90 | 0.98 | 0.97 |
| GraphSage | 0.71 | 0.65 | 0.62 | 0.64 | 0.87 |
| ComplEX | 0.84 | 0.76 | 0.74 | 0.95 | 0.99 |
| DistMult | 0.93 | 0.93 | 0.92 | 0.95 | 0.98 |
| TransE | 0.88 | 0.87 | 0.87 | 0.95 | 0.95 |
| **rd-explainer** | **0.95** | **0.95** | **0.95** | **0.99** | **0.99** |

### 4.4. Drug predictions validation based on the scientific literature

We also evaluated the prediction performance based on the capacity of our method to discover marketed drugs already reported being used for a new phenotype. First, we listed for each of the 7 selected phenotypes the three drugs with the highest scores. Because the objective is to find new indications for drugs; if any of the reported drugs already appears in the graph as a treatment for the targeted symptom, this drug will be skipped and the next one with the highest score will be selected. For example, if aprindine is selected as the drug with the highest score to treat arrhythmia, but the relation ’aprindine is a substance that treats arrhythmia’ is already present in our graph, aprindine won’t be reported as a possible drug candidate.

For each possible drug candidate, a literature search was carried out to find preliminary evidence if that drug had already been used to treat the symptom. If the drug was contraindicated to treat the symptom (or if it could cause the symptom) it was also annotated. Results regarding each drug candidate obtained using KG A can be found in Table S9. Additionally, Table 6 summarizes the amount of drugs (in percentage) that contained supporting evidence, contraindication evidence or no evidence at all. We found that only a fifth of the drug candidates had supporting evidence in the literature, and that the vast majority of the candidates (65.43%) did not have any evidence at all. There is a small percentage of them that are actually contraindicated to treat the targeted symptom/phenotype. Finally, the amount of supporting/contraindicating evidence can be found summarized in Table S7.

**Table 6.** Percentage of drugs containing supporting evidence, contraindication evidence or no evidence at all for both Graph A and B.

| *Property* | **KG A** | **KG B** |
| --- | --- | --- |
| *Supporting Evidence* | 20.99 % | 27.16 % |
| *Contraindication Evidence* | 13.58 % | 14.82 % |
| *No Evidence* | 65.43 % | 58.02 % |

The same approach was followed in the case of KG B. Information regarding the drug candidates for each symptom (as well as the supporting evidence) can be found also in Table S10. Additionally, the percentage of drugs with supporting evidence, contraindication evidence or no evidence at all can be seen in Table 6. In this case, the number of drug candidates with evidence has increased with respect to the drug candidates obtained with KG A (27% in B vs 21% in A), and the number of drug candidates with no evidence has been reduced (58% in B vs 65% in A). The number of drug candidates with contraindications remains almost the same (13% in A vs 14% in B).

### 4.5. Evaluating drug repurposing explanations as semantic graphs

Evaluating an explanation is a tough task and many different benchmarks are recently appearing to evaluate them [39]. In this work, we followed two different approaches to evaluate the explanations: a more subjective one, where the explanation was evaluated with our own biological knowledge; and a more objective one, where a manual literature search and curation was performed to check if the suggested explanation has already been reported. We selected 7 phenotypes (muscular dystrophy, respiratory insufficiency, arrhythmia, dilated cardiomyopathy, congestive heart failure, progressive muscle weakness and cognitive impairment) and their top 3 predictions, then explanations were produced from the models trained on both KGs. The selection of these phenotypes aimed to cover the diverse systems affected by the disease. Each explanation was analyzed and, if possible, compared to the one that was found in the literature.

Explanations were classified into complete and incomplete explanations. Complete explanations are those that show a connection (path) between the drug candidate and the targeted symptom/phenotype (Figure S3). They are considered complete as they allow for an easy human-understandable interpretation. On the other hand, incomplete explanations are those where the explanation is composed of two separated clusters (one for the drug and one for the phenotype) (Figure S4) or by a unique cluster where either the drug or the phenotype is missing (Figure S5).

The global analysis of the completeness of explanations generated can be seen in Table 7 (amount of complete and incomplete explanations in each type of supporting evidence) and Table 8 (amount of supporting evidence in each type of explanations). This analysis was performed taking into account the explanations from both graphs. As it can be seen in Table 7, in total the same number of complete and incomplete explanations was obtained (21 each). However, when looking at each category separately, it is seen that when there is evidence GNNExplainer tends to produce complete explanations (68 %), and conversely when there is no supporting evidence or when the drug is contraindicated the resulting explanation is usually incomplete (62 % and 70 %, respectively). As it can be seen in Table 8, when a complete explanation is created, almost 2/3 of the time the explanation contains supporting evidence (62 %); while when the explanation is incomplete, only 1/4 of the times it contains supporting evidence (28 %).

**Table 7.** Number and percentage of complete and incomplete explanations in each evidence type.

|  | Complete Explanations | Percentage Complete Explanations | Incomplete Explanations | Percentage Incomplete Explanations |
| --- | --- | --- | --- | --- |
| *Supporting Evidence* | 13 | 68 % | 6 | 32 % |
| *Contraindication Evidence* | 3 | 30 % | 7 | 70 % |
| *No Evidence* | 5 | 38 % | 8 | 62 % |
| *Total* | 21 | 50 % | 21 | 50 % |

**Table 8.**
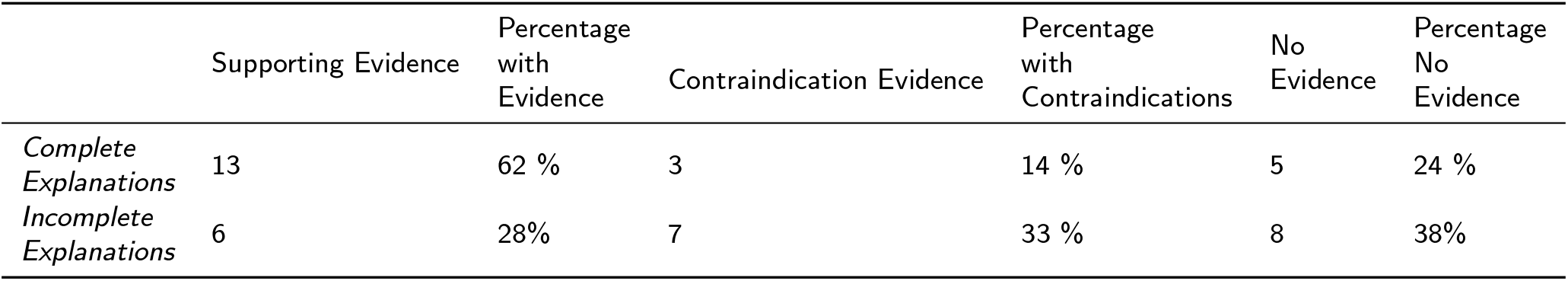
Number and percentage of explanations with no evidence, with supporting evidence and with contraindications in each type of explanation.

An additional analysis was performed, this time considering each graph separately. This can be seen in Table 9 and Table S7. There is a clear difference between the explanations obtained in graph A and B. Firstly, KG A explanations are more likely to be complete (72 % in A vs 28 % in B), while KG B produces more incomplete explanations (72 % in B vs 28 % in A) (Table 9).

**Table 9.** Number and percentage of complete and incomplete explanations in each evidence type and in each graph.

|  | Evidence Type | Complete Explanations |  | Incomplete Explanations |  |
| --- | --- | --- | --- | --- | --- |
|  |  | Number | Percentage | Number | Percentage |
| KG A | Supporting Evidence | 9 | 100% | 0 | 0% |
|  | Contraindication Evidence | 1 | 17% | 5 | 83% |
|  | No Evidence | 5 | 83% | 1 | 17% |
|  | **Total** | **15** | **72%** | **6** | **28%** |
| KG B | Supporting Evidence | 4 | 40% | 6 | 60% |
|  | Contraindication Evidence | 2 | 50% | 2 | 50% |
|  | No Evidence | 0 | 0% | 7 | 100% |
|  | **Total** | **6** | **28%** | **15** | **72%** |

An example of an explanation produced by rd-explainer can be seen in Figure 5. This explanation is classified into complete and suggests why Doxorubicin should be considered for treating respiratory insufficiency; as it is a drug that targets CHRM1 a gene that interacts with DAG1, which causes the disease. Throughout this section explanations have been classified into complete and incomplete. However, an explanation being complete does not make it a good explanation. This way, for example, an explanation of the type ‘Drug A targets Gene B, Gene B interacts with Gene C, and Gene C causes Disease D’ can make biological sense such as in Figure 5. On the other hand, an explanation of the type ‘Drug A treats Disease B, Disease B is caused by Gene C, Gene C causes Disease D’ does not make full biological sense (Drug A could treat Disease B by targeting a gene other than Gene C; this way, the same treatment could not be applied for Disease D). This is in fact what is observed in Figure S6, where disopyramide is said to treat muscular dystrophy following the next explanation: disopyramide treats urinary incontinence, affectation in DMD gene can cause urinary incontinence, and DMD gene has as phenotype muscular dystrophy. In this case, a person may have urinary incontinence for several reasons, and disopyramide may be able to treat one of them, but not necessarily the one caused by affectation in DMD gene.

**Figure 5:**
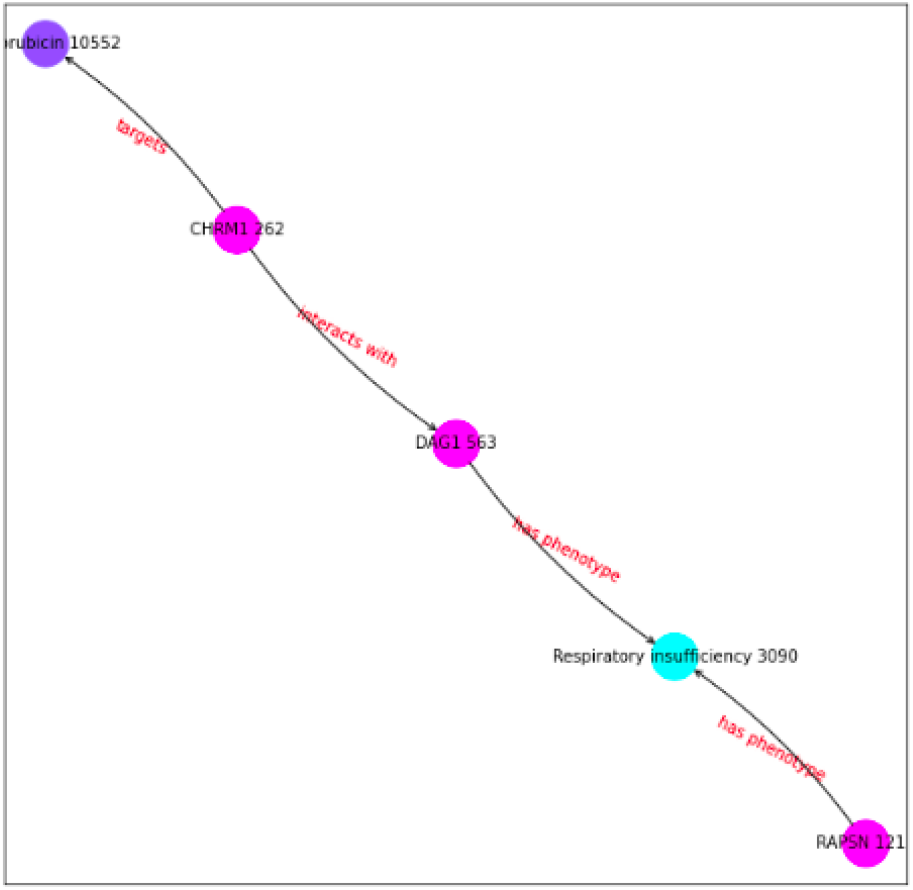
Explanation of drug candidate Doxorubicin as possible treatment for Respiratory Insufficiency. Classified as complete explanation.

The objective evaluation is undoubtedly more unbiased and equitable. Nonetheless, subjective evaluations are also significant since there are drug-phenotype interactions that are not fully understood (specially when a certain drug is producing an undesired side effect), and so they are not well established in the literature. But, analyzing the proposed explanations based on expert domain knowledge might shed light on the interaction and help to formulate a hypothesis that can be clearly designed to be tested in the wet laboratory. After applying the objective evaluation only one explanation (levosimendan - progressive muscle weakness) was found to have supporting evidence (where levosimendan treats the disease by increasing the troponin C affinity for calcium), and two links’ explanations (doxorubicin - respiratory insufficiency and sorafenib - respiratory insufficiency) contained unclear interactions (both were of type contraindications). The results after applying this evaluation can be found summarised in the Table S6. Regarding the subjective evaluations, 17 out of 21 explanations were found to be good explanations (they were in accordance with biological reasoning) such as the one illustrated by doxorubicin - respiratory insufficiency in Figure 5; and 4 were considered bad explanations (they made no biological sense), the previously mentioned disopyramide - muscular dystrophy in Figure S6, and the explanations in Figures S7, S8 and S9.

### 4.6. Generalizability of rd-explainer tested on other case studies

To show that this method can be extended to other rare diseases it was also tested in Alzheimer’s Disease (AD) and Amyotrophic Lateral Sclerosis (ALS) type 1. Despite Alzheimer Disease not being a rare disease, there are different types of Alzheimer with very little prevalence. This way, for the Alzheimer’s knowledge graph we used the general disease (MONDO:0004975) and all its causal genes that were present in Monarch (APP (HGNC:620), APOE (HGNC:613), PSEN1 (HGNC:9501) and PSEN2 (HGNC:9509)) as seeds. The final result would be a knowledge graph that specializes in Alzheimer diseases and that we can use to focus on the symptoms of the rare types of the disease. For the ALS type 1 knowledge graph we used the seed for the disease (MONDO:0007103) and the causal gene according to Monarch (SOD1 (HGNC:11179)). Table 10 shows the GNN performance in both diseases, showing once more a high AUROC and AUPRC for these diseases.

**Table 10.**
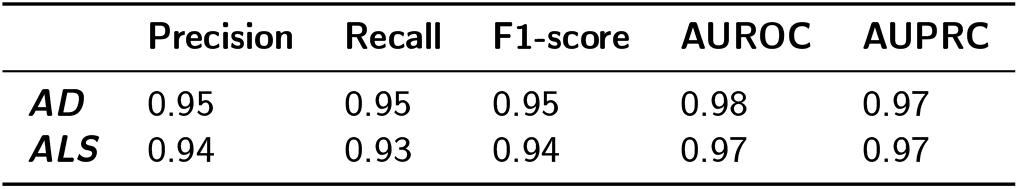
Table showing different performance metrics tested in AD and ALS.

Next, the same approach that was followed for DMD was followed for both diseases: for each symptom we analysed the three drug candidates with the highest score (this drug candidates should not appear in the knowledge graph); then a literature search was performed to check if the drug candidates had been reported by the scientific community. The complete list of phenotypes as well as the drug candidates and scores for each phenotype can be found in Tables S11 and S12. This tables also contain whether the drug candidates had supporting evidence in the literature.

Among the predictions, it is worth mentioning Pexidartinib, a drug candidate that was proposed by the model to treat memory impairment in AD and that is currently undergoing a clinical trial as a drug that could be potentially beneficial to treat the disease [40].

## 5. Discussion

We integrated disease-specific knowledge graphs in combination with GNN and XAI for interpretable drug repurposing. We found that state-of-the-art XAI methods based on GNNs support *in silico* predictions of candidate repurposable drugs for rare diseases by providing interpretable reasoning paths of mechanism of action. We developed *rd-explainer*, a method to perform computational drug repurposing specifically for rare diseases. It utilizes cutting-edge deep learning methods such as edge2vec and GNNs and provides drug-symptom/phenotype predictions with high performance scores, and utilizes a modified version of GNNExplainer to provide explanations as semantic graphs for the interpretability of the results. We also found that these explanations have different levels of usefulness to generate testable hypotheses: paths linking drug and phenotype nodes are more understandable versus isolated clusters since they are similar to human reasoning; adding semantics to relations adds biological meaning to help to formulate a hypothesis and design the experiment in the laboratory; and providing clear semantic graphs by removing relations that are not contributors in the learning process. We tested the generalizability of our method executing it on two additional diseases: ALS and AD. ALS type 1 was selected to test the pipeline in another monogenic disease with fewer information available. AD was selected as it is a common disease with rare subtypes that can be caused by several genes, and we wanted to test the pipeline in a polygenic and multifactorial disease. We demonstrated that our pipeline performs well on mono- and polygenic rare diseases.

rd-explainer is a researcher-centered drug repurposing method that has been demonstrated as an innovative AI based method for rare disease drug research. rd-explainer’s main advantage is its interpretability. The main motivation of this study was to provide explanations underlying AI predictions. rd-explainer provides explanations as semantic graphs, a type of explanation that resembles to human reasoning. This is in line with current research on user-centric XAI [41]. Not only does this have the high value to support rare disease researchers to formulate evidence-based hypotheses testable in the wet laboratory (and reduce cost, time and risk), but to gain new disease knowledge and speed up robust drug research. Our approach was to use state-of-the-art AI and XAI methods used in drug repurposing such as knowledge graphs to naturally represent known associations among biological entities with expressive semantics and supporting curated evidence, graph learning, and graph based XAI methods. The advance in the rare disease field is that we provide interpretable predictions thanks to a pipeline that it seamlessly integrates a graph learning model with an explainer, combining results of both model performance and explanation accuracy to mitigate the black-box problem and foster XAI adoption in the field [42]. BioKnowledge Reviewer tool provides rare disease specific knowledge graphs for disease biology data collection by means of the Monarch knowledge base API [29]. We argue that a tool or approach that can collect associations from a virtual, federated knowledge graph via APIs could extend this feature to any biomedical associations such as for drug data collection, and improve data and knowledge driven research. Another great advantage of the rd-explainer method is its modular implementation; this means that different parts of the workflow (data, features, GNN and explanations) can be independently modified and the pipeline can still be run. For example, if one is interested in using another node feature embedding algorithm instead of edge2vec, one can just modify that component of the pipeline and still run the rest of the workflow.

Our results showed that rd-explainer is a highly performant graph ML based drug repurposing method. Our method builds rare disease-specific models trained on newly generated KG for the disease of focus and enriched with data for the prediction task. In comparison with state-of-the-art AI-based drug repurposing approaches, rd-explainer demonstrates outstanding performance. Throughout this paper, we have compared rd-explainer with various AI methods that employ different techniques for their predictions, including GNNs such as GraphSAGE, random walk embeddings like edge2vec, and geometric embeddings using models like ComplEX, DistMult, and TransE. By combining random walk models (edge2vec) with GNNs (GraphSAGE), rd-explainer achieves superior results in the link prediction task. Notably, edge2vec outperforms GraphSAGE, suggesting that the exceptional performance of rd-explainer is primarily attributed to the random walk model, with the GNN providing an additional performance boost. This level of performance rivals other models developed for drug repurposing, such as deepDR (AUROC = 0.908) [43]. Although there are benchmarks and frameworks to evaluate the performance of GNNs [44, 45, 46, 47, 48, 49] to the best of our knowledge there is no a standard for drug repurposing, and makes it challenging to directly compare rd-explainer to other methods due to one of its key features: the creation of high-quality disease specific knowledge graphs. These knowledge graphs are enriched with data from a wide array of sources including domain expert knowledge via the seed nodes, and curated known relations among genes, anatomical structures, biological processes and diseases not only from humans, but also importantly numerous other species to fill the lack of molecular knowledge. This comprehensive approach significantly boosts the graph’s richness and diversity, making it a valuable resource for tackling rare diseases, which often suffer from limited research attention. By maximizing the information available, rd-explainer enhances our ability to identify potential treatments for these understudied conditions and, ultimately, enable more effective and faster translation. Conversely, Huang et al. recently proposed a clinician-centered drug repurposing foundation model pretrained on a medical KG composed of 17.000 diseases and transfer learning by disease mechanism similarity [50]. It would be interesting to combine both approaches and investigate the effect of extending our KGs with similar disease networks from well-known diseases.

Our new predictions are valid drug candidates since they are consistent with recent findings in the literature. We demonstrated that rd-explainer can provide new interesting drug-phenotype predictions. For instance, Sunitinib, one of the drugs that appear to be a good candidate to treat the symptoms of the disease according to both models (using KG A and KG B), has been considered as a good drug candidate to treat DMD and in 2019 appeared to be in preclinical trials [51]. This drug belongs to the group of tyrosine kinase inhibitors, and many other drugs that belong to this category have been proposed by our model (Fedratinib, Sorafenib, Bosutinib, Ruxolitinib and Midostaurin). Similarly, Mezlocillin, an antibiotic used to treat gram-negative bacterial infections, has also been proposed by our model; while Gentamicin, another gram-negative antibiotic, was in 2019 in clinical trials to treat DMD [51]. This way, despite not producing drugs candidates that are undergoing a clinical trial or a treating the disease, it produces drug candidates that participate in similar biological processes (ie. tyrosine kinases inhibitor, gram negative antibiotics)

Importantly, explanations for hypothesis generation may enable to move towards *lab-in-the-loop* framework. With respect to the interpretability and utility of explanations, one of the 21 examined explanations was supported by evidence in the literature. Nonetheless, this does not mean that the explanations are useless. A good example of this would be the explanation for the Methylprednisolone-Muscular Dystrophy link (Figure S10). The explanation is simple: ‘Methyl-prednisolone treats DMD, DMD has Muscular Dystrophy as phenotype; thus methylprednisolone can treat Muscular Dystrophy’. In this case the explanation does not contain supporting evidence but the explanation still makes sense. In the literature, methylprednisolone is said to be a good candidate to treat muscular dystrophies because it interacts with the glucocorticoid receptor and this leads to the activation of anti-inflammatory signaling and the inhibition of proinflammatory signaling [52]. The explanation proposed by rd-explainer doesn’t provide the underlying causative mechanism that relates methylprednisolone and muscular dystrophy, but a researcher can still be able to see that muscular dystrophies and methylprednisolone are interrelated. This illustrates how even though an explanation may lack comprehensive supporting evidence, it can still provide valuable directional cues for further more precise investigation. Another important aspect is that rare disease findings in the lab can be introduced back in the knowledge graph to update and improve the disease specific AI model for continual learning and enabling precise experimental design. Besides, this synergy fosters collaboration between computational and wet lab researchers to increase efficiency for disease specific drug research [29].

Finally, we found that knowledge graph topology has an impact on explainability. It was also seen that KG A usually produces more complete explanations, while in KG B incomplete explanations appear to be more numerous. This could happen due to the difference in the graph structure itself: graph A has a smaller clustering coefficient than graph B (see Section 4.1), which leads to more edges being present in the subgraphs produced by GNNExplainer. This way, because the 15th edges with the highest scores are selected, it is more likely to find a path between drug and phenotype in KG A than in B. Another interesting difference is that explanations generated with KG A tend to have a higher ’sensitivity’, while explanations generated with KG B tend to have a higher ’specificity’. When an incomplete explanation is produced using graph A it is very unlikely that the explanation will contain supporting evidence (0 explanations were found to have evidence if the explanation was incomplete in KG A). Similarly, when a complete explanation is produced in KG B, it is very likely that the explanations have supporting evidence or contraindication evidence (67% of complete explanations had supporting evidence and 33% of complete explanations had contraindication evidence). For this reason, if one remains skeptical about the explanations themselves, this quality of the explanations might be used as filter/validation. For example, if an incomplete explanation is obtained with KG A, it is unlikely that it is trustworthy (none of the incomplete explanations had supporting evidence). Similarly, if a complete explanation is obtained using KG B, it is likely that there is some interaction between the drug and the phenotype (all of the complete explanations generated with graph B had either supporting or contraindication evidence). Our findings are aligned with recent studies where the influence of clustering coefficient and topology has been observed in embedding-based predictions [53, 54], here we extend these observations to its impact on graph-based explanations.

### Limitations and future directions

An important limitation of this study is that we only utilize one XAI method, which is not model agnostic. XAI is a hot research topic in the AI field, where new and more sophisticated methods are frequently published [55]. It would be good to extend our study to other XAI types to check how applicable they are given the unique characteristics of rare diseases, including limited annotated data, lack of knowledge of pertinent entity relations, and lack of a gold drug-phenotype standard. Another important limitation is the lack of standard benchmarking and metrics to systematically evaluate explainers and explanations. Currently, there are some initial efforts going in this direction [56, 57, 34, 58, 59, 60], but there is still a lack of a common standard [61]. The known reproducibility issue of our explainer [34] that may imply that the explanations are different each time it is used, may reduce the confidence and reliance on the explanations. We did several experiments to try and bring consistency to explanations; for example, executing GNNExplainer several times and using the mean mask as the final mask or increasing the number of epochs. However, this still did not solve the issue. This experience makes us strongly recommend to work on the standard evaluation of explanations by the XAI community to foster trust on the application of AI in bioinformatics and biomedicine. Additionally, many times the explanation would consist in a subgraph where the two targeted nodes would be disconnected from each other, which might bring confusion and could be seen as a ‘bad’ explanation. Therefore, work towards methods that prioritize or focus on providing just connecting paths such as metapath based ones [62, 63, 64, 65, 66] and on improving path visualisation for user interpretation [67, 68, 69] is arguably recommended. Finally, while we focused primarily on integrating a graph ML model with an explainer, a clear line of research will be to work on interpretability and reproducibility of explanations in the context of the drug repurposing task. The reproducibility/incosistency could be affected by the size and complexity of our data. This inconsistency could make the users of this pipeline skeptical about its explanations and for this reason more investigation should be done in this element of the pipeline to make it a more robust model. To improve this, ontologies could be incorporated into the knowledge graph to increase the quality and interpretability of our data. Ontologies help to standardize data into the shared meaning by a community enhancing thus interpretability by domain users. Importantly, the formal description of knowledge embedded in ontologies can be leveraged for data consistency checking, and for inference to add implicit knowledge into the graph [70]. Nonetheless, knowledge graph and ontology changes pose a great interoperability challenge to the community to keep up downstream bioinformatics and data science workflows and analyses [71, 72]. Finally, it would make our work more ’FAIR’ [73], i.e., not only understandable by humans, but also by machines, by providing our drug repurposing for DMD KG from a FAIR data point [74], and rd-explainer from workflowHub [75].

## 6. Conclusion

We present the application of explainable AI on state-of-the-art computational drug repurposing for rare diseases. Our knowledge graph based deep learning method provides human understandable explanations for the phenotype-drug link prediction and we demonstrated that graph XAI can be applied to rare diseases. The *rd-explainer* method provides an innovative approach that can maximize the available disease-specific knowledge and generate valuable predictions with its explanations. Our model has proven to obtain high evaluation scores, providing drug candidates that are often supported by evidence. The key contribution of our study is that our pipeline gives possible explanations in the form of semantic graphs that may help rare disease researchers to make informed decisions to experimentally validate deep learning model predictions. However, we detected that data topology affects explanations, highlighting the importance of investigating further how best represent graphical knowledge for model performance and explanation accuracy. *rd-explainer* can be extended to other rare diseases and provide computer-aided guidance for biologists and accelerate translational research. Finally, future studies should advance our understanding of the necessary standard mechanism to evaluate explainability to foster adoption from domain experts and to mitigate the black-box problem of trust on AI, especially for biomedicine where decisions can have an important impact on people’s lives.

## Supporting information

Supplementary section

## 7. Code availability

The code to run the repurposing pipeline is available at https://github.com/PPerdomoQ/rare-disease-explainer.

## Footnotes

1 https://github.com/RoyZhengGao/edge2vec

2 DrugCentral, *Download site*, accessed March 2022, https://unmtid-dbs.net/download/DrugCentral/2021_09_01/drug.target.interaction.tsv.gz

3 Therapeutic Target Database, *Download site*, accessed March 2022,https://idrblab.net/ttd/sites/default/files/ttd_database/P1-05-Drug_disease.txt

