## Supplementary section for "Knowledge Graphs and Explainable AI for Drug Repurposing on Rare Diseases"

### A. Supplementary section

#### A.1. Knowledge Graph seeds

**Table S1**
Table containing seeds used to build KG A.

| *Seed Name* | Seed ID |
|---|---|
| *DMD* | HGNC:2928 |
| *DMD* | MONDO:0010679 |

**Table S2**
Table containing seeds used to build KG B.

| *Seed Name* | Seed ID |
|---|---|
| *DMD* | HGNC:2928 |
| *DMD* | MONDO:0010679 |
| *Hypotonia* | HP:0001252 |
| *Specific learning disability* | HP:0001328 |
| *Arrhythmia* | HP:0011675 |
| *Congestive heart failure* | HP:0001635 |
| *Dilated cardiomyopathy* | HP:0001644 |
| *Calf muscle hypertrophy* | HP:0008981 |
| *Motor delay* | HP:0001270 |
| *Muscular dystrophy* | HP:0003560 |
| *Delayed speech and language development* | HP:0000750 |
| *Hypoventilation* | HP:0002791 |
| *Intellectual disability, mild* | HP:0001256 |
| *Hyporeflexia* | HP:0001265 |
| *Cognitive impairment* | HP:0100543 |
| *Proximal muscle weakness* | HP:0003701 |
| *Abnormal EKG* | HP:0003115 |
| *Calf muscle pseudohypertrophy* | HP:0003707 |
| *Cardiomyopathy* | HP:0001638 |
| *Flexion contracture* | HP:0001371 |
| *Elevated circulating creatine kinase concentration* | HP:0003236 |
| *Global developmental delay* | HP:0001263 |
| *Skeletal muscle atrophy* | HP:0003202 |
| *Respiratory insufficiency* | HP:0002093 |
| *Waddling gait* | HP:0002515 |
| *Gowers sign* | HP:0003391 |
| *Generalized hypotonia* | HP:0001290 |
| *Progressive muscle weakness* | HP:0003323 |
| *Scoliosis* | HP:0002650 |
| *Hyperlordosis* | HP:0003307 |

#### A.2. Number of edge types

**Table S3**
Number and percentage of edge types in KG A.

| Edge Type | Count | Percentage |
|---|---|---|
| *in 1 to 1 orthology relationship with* | 35650 | 37.96% |
| *in orthology relationship with* | 25242 | 26.88% |
| *has phenotype* | 15730 | 16.75% |
| *interacts with* | 9824 | 10.46% |
| *is part of* | 1465 | 1.56% |
| *has affected feature* | 1101 | 1.17% |
| *expressed in* | 1079 | 1.14% |
| *enables* | 983 | 1.04% |
| *pathogenic for condition* | 976 | 1.03% |
| *targets* | 518 | 0.55% |
| *involved in* | 432 | 0.46% |
| *likely pathogenic for condition* | 182 | 0.19% |
| *contributes to condition* | 171 | 0.18% |
| *has role in modeling* | 134 | 0.14% |
| *is allele of* | 96 | 0.10% |
| *is substance that treats* | 86 | 0.09% |
| *colocalizes with* | 84 | 0.09% |
| *source* | 29 | 0.03% |
| *is causal germline mutation in* | 16 | 0.02% |
| *has genotype* | 7 | 0.01% |
| *contributes to* | 5 | 0.01% |
| *causes condition* | 3 | 0.003% |
| *is marker for* | 1 | 0.001% |
| *is causal germline mutation partially giving rise to* | 1 | 0.001% |

**Table S4**
Number and percentage of edge types in the KG B.

| Edge Type | Count | Percentage |
|---|---|---|
| *has phenotype* | 836138 | 42.13% |
| *in 1 to 1 orthology relationship with* | 520547 | 23.23% |
| *in orthology relationship with* | 333288 | 16.79% |
| *interacts with* | 226174 | 11.40% |
| *expressed in* | 14589 | 0.74% |
| *is part of* | 9427 | 0.47% |
| *colocalizes with* | 8112 | 0.41% |
| *involved in* | 7790 | 0.39% |
| *enables* | 7053 | 0.36% |
| *targets* | 5070 | 0.26% |
| *has role in modeling* | 3449 | 0.17% |
| *causes condition* | 2479 | 0.12% |
| *contributes to condition* | 2203 | 0.11% |
| *is allele of* | 1167 | 0.06% |
| *has affected feature* | 1137 | 0.06% |
| *pathogenic for condition* | 1024 | 0.05% |
| *is causal germline mutation in* | 900 | 0.04% |
| *is substance that treats* | 599 | 0.03% |
| *contributes to* | 198 | 0.01% |
| *likely pathogenic for condition* | 185 | 0.01% |
| *is causal loss of function germline mutation of in* | 179 | 0.01% |
| *is reference allele of* | 130 | 0.01% |
| *is marker for* | 97 | 0.005% |
| *has genotype* | 67 | 0.003% |
| *is causal susceptibility factor for* | 42 | 0.002% |
| *source* | 32 | 0.002% |
| *is causal somatic mutation in* | 16 | 0.001% |
| *is causal gain of function germline mutation of in* | 15 | 0.001% |
| *is causal germline mutation partially giving rise to* | 12 | 0.001% |

##### A.3. GNNExplainer algorithm

```
Input: GNN, NodeIdx1, NodeIdx2, G
Output: G_{s,m}, Mask
Emb = GNN(G) // Obtain embeddings
InitialPred = Emb[NodeIdx1] · Emb[NodeIdx2] // Get initial prediction
G_s = Subgraph(G, NodeIndex1, NodeIndex2) // Obtain subgraph
Mask = InitializeMask(G_s) // Initialize Mask
for Epoch in Epochs do
    G_{s,m} = ApplyMask(G_s, Mask) // Apply Mask to subgraph
    NewEmb = GNN(G_{s,m}) // Get new embeddings
    NewPred = NewEmb[NodeIdx1] · NewEmb[NodeIde2] // Get new prediction
    Loss = GetLoss(InitialPred, NewPred) // Calculate loss
    Mask = Backpropagate(Mask, Loss) // Backpropagate loss
end
return G_{s,m}, Mask
```

**Algorithm 1:** GNNExplainer Link Prediction Pseudocode. *GNN* stands for the trained GNN model. *G* stands for the Graph.

#### A.4. List of hyperparameters

**Table S5**
Table showing the different options of hyperparameters that were tested as well as their optimal values.

| ***Process*** | **Hyperparameter** | **Options** | **Optimal Value** |
|---|---|---|---|
| *edge2vec* | Number of walks | 2, 4, 6 | 2 |
| | Walk Length | 3, 5, 7 | 7 |
| | Embedding Dimension | 32, 64, 128 | 32 |
| | Edge Direction | Undirected, Directed | Directed |
| | p | 0.5, 0.7, 1 | 0.7 |
| | q | 0.5, 0.7, 1 | 1 |
| | Epochs | 5, 10 | 10 |
| *GNN* | Hidden Dimension | 64, 128, 256 | 256 |
| | Output Dimension | 64, 128, 256 | 64 |
| | Layers | 2, 4, 6 | 2 |
| | Aggregation Function | mean, sum | mean |
| | Dropout | 0, 0.1, 0.2 | 0.2 |
| | Learning Rate | 0.001 - 0.1 | 0.07 |
| | Epochs | 100, 150, 200 | 150 |

#### A.5. Visualization of explanations

To visualize the resulting explanations, a custom visualization function was developed to represent explanations as more human readable and semantic graphs and, thus, improving the one provided by Pytorch Geometric [76]. In the first place, the possibility of visualizing the edge types has been incorporated. Additionally, in this new formula several customizable parameters have been added. Now, it is possible to only visualize the active edges of the explanation, removing non-important edges. This will allow for clearer visualization of the subgraph. Figure S1 shows how an explanation is modified after applying this option. Finally, it is also possible to remove unconnected clusters from the explanations. This way, if an explanation is formed by several clusters, there is the possibility of just viewing the ones that contain the drug candidate and the targeted phenotype. Figure S2 shows how the explanation is modified after applying this filter.

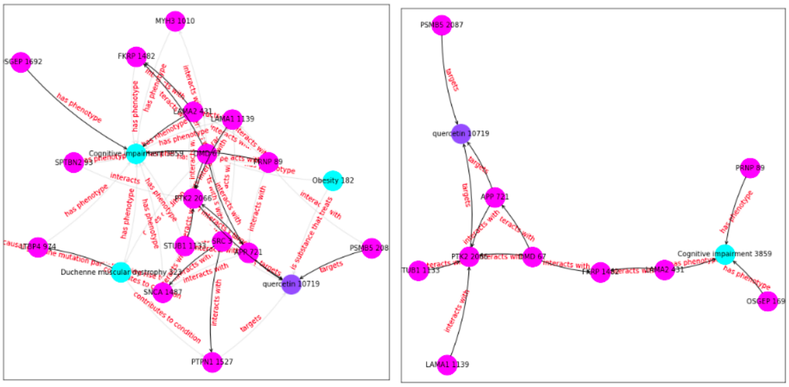


**Figure S1:** Explanation after removing non-important edges. Left: Explanation keeping all the edges. Right: Explanation removing non-important edges.

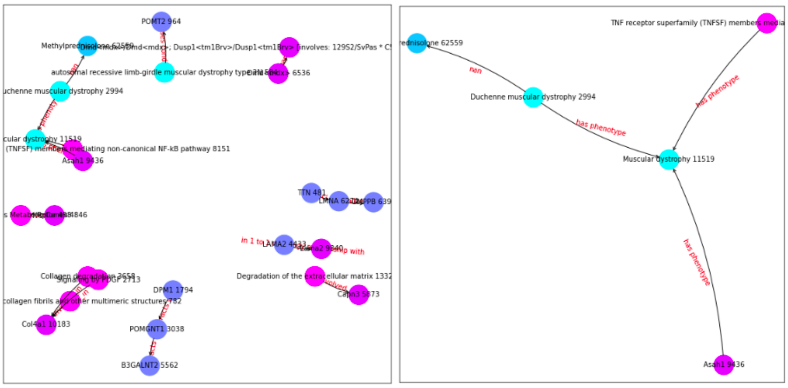


**Figure S2:** Explanation after removing unconnected clusters. Left: Explanation keeping all the clusters. Right: Explanation removing additional clusters.

#### A.6. Complete/Incomplete explanation Example

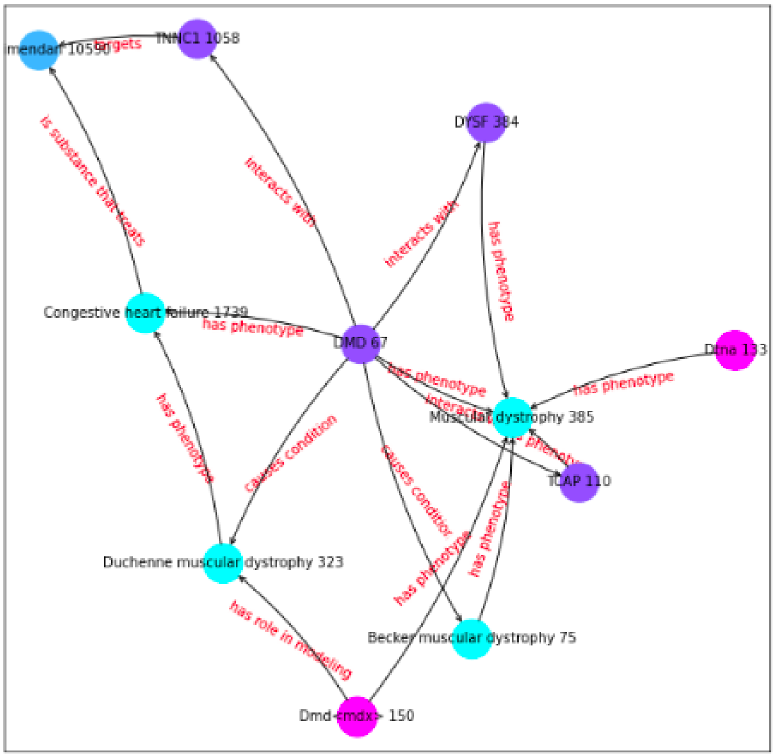


**Figure S3:** Explanation of drug candidate Levosimendan as possible treatment for Muscular Dystrophy. Classified as complete explanation.

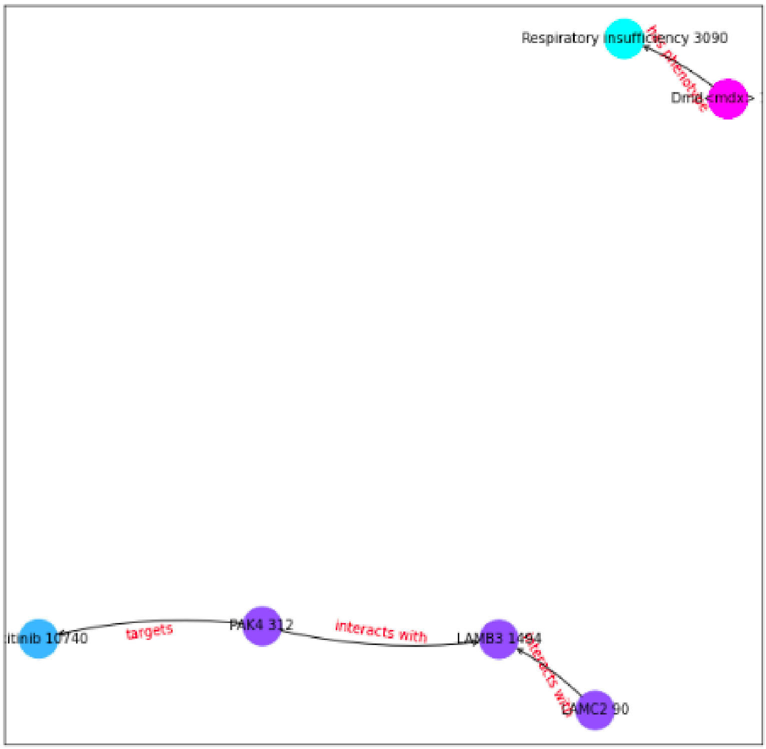


**Figure S4:** Explanation of drug candidate Axitinib as possible treatment for Respiratory Insufficiency. Classified as incomplete explanation.

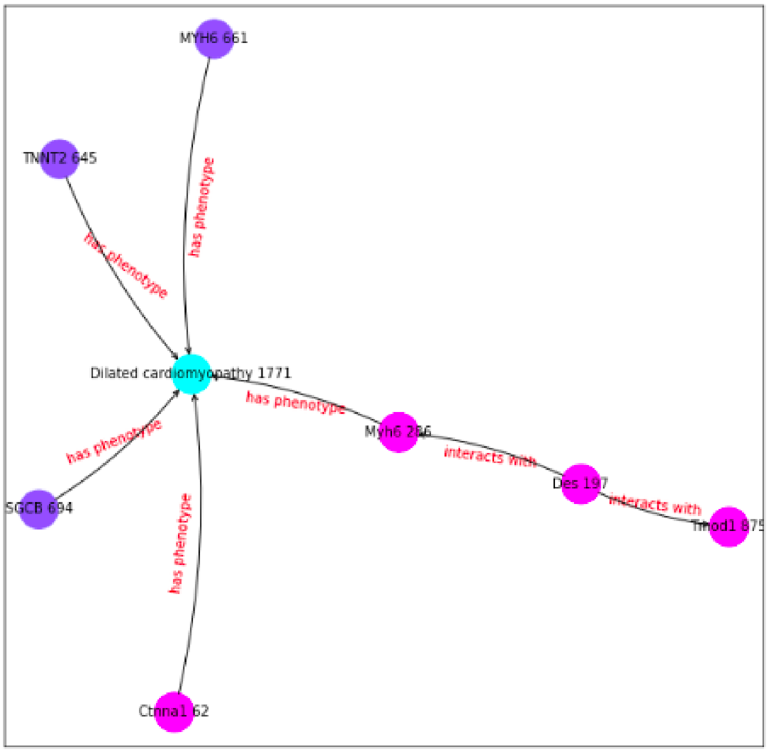


**Figure S5:** Explanation of drug candidate Entrectinib as possible treatment for Dilated cardiomyopathy. Classified as incomplete explanation.

#### A.7. Evaluation of explanations

**Table S6**
Table showing the amount of times each drug appears as one of the top 3 drug candidates with highest score treat one of the 27 symptoms. It is also shown the amount of supporting evidence and contraindication evidence for each drug. This information was obtained using Graph A.

| Drug | Appearances | Percentage | With Evidence | With Contraindications |
|---|---|---|---|---|
| *Entrectinib* | 25 | 92.59 % | 0 | 8 |
| *Axitinib* | 19 | 70.37 % | 1 | 1 |
| *Nintedanib* | 12 | 44.44 % | 2 | 0 |
| *Levosimendan* | 7 | 25.92 % | 6 | 0 |
| *Disopyramide* | 6 | 22.22 % | 2 | 0 |
| *Doxorubicin* | 2 | 7.40 % | 0 | 2 |
| *Aprindine* | 2 | 7.40 % | 2 | 0 |
| *Amiodarone* | 1 | 3.70 % | 1 | 0 |
| *Acepromazine* | 1 | 3.70 % | 0 | 0 |
| *Mezlocillin* | 1 | 3.70 % | 0 | 0 |
| *Sunitinib* | 1 | 3.70 % | 0 | 0 |
| *Fedratinib* | 1 | 3.70 % | 0 | 0 |
| *Carvedilol* | 1 | 3.70 % | 1 | 0 |
| *Queracetin* | 1 | 3.70 % | 1 | 0 |

**Table S7**
Table showing the number and percentage of explanations with no evidence, with supporting evidence, and with contraindications for each type of explanation and each graph.

| | | With Evidence | Percentage With Evidence | With Contraindications | Percentage With Contraindications | No Evidence | Percentage No Evidence |
|---|---|---|---|---|---|---|---|
| *KG A* | *Complete Explanations* | 9 | 60% | 1 | 7% | 5 | 33% |
| | *Incomplete Explanations* | 0 | 0% | 5 | 83% | 1 | 17% |
| *KG B (Large)* | *Complete Explanations* | 4 | 67% | 2 | 33% | 0 | 0% |
| | *Incomplete Explanations* | 6 | 40% | 2 | 13% | 7 | 47% |

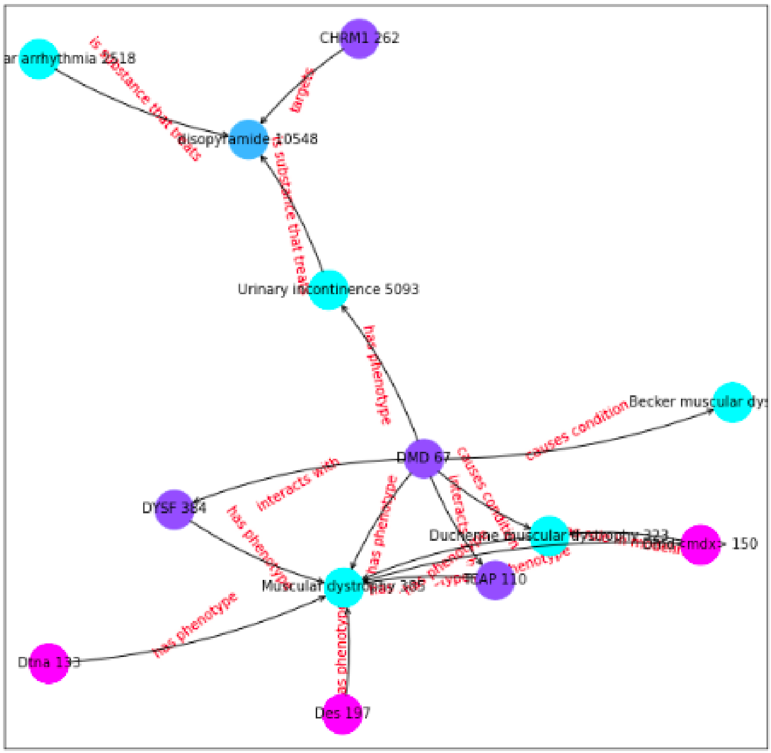


**Figure S6:** Explanation of drug candidate Disopyramide as possible treatment for Muscular Dystrophy. Classified as complete explanation.

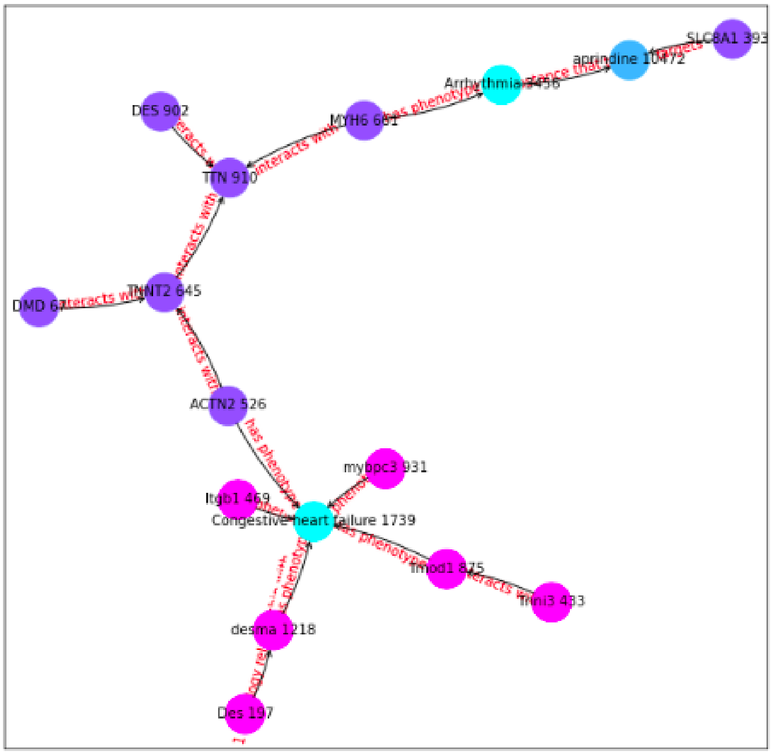


**Figure S7:** Explanation of drug candidate Aprindine as possible treatment for Congestive heart failure. Classified as complete explanation.

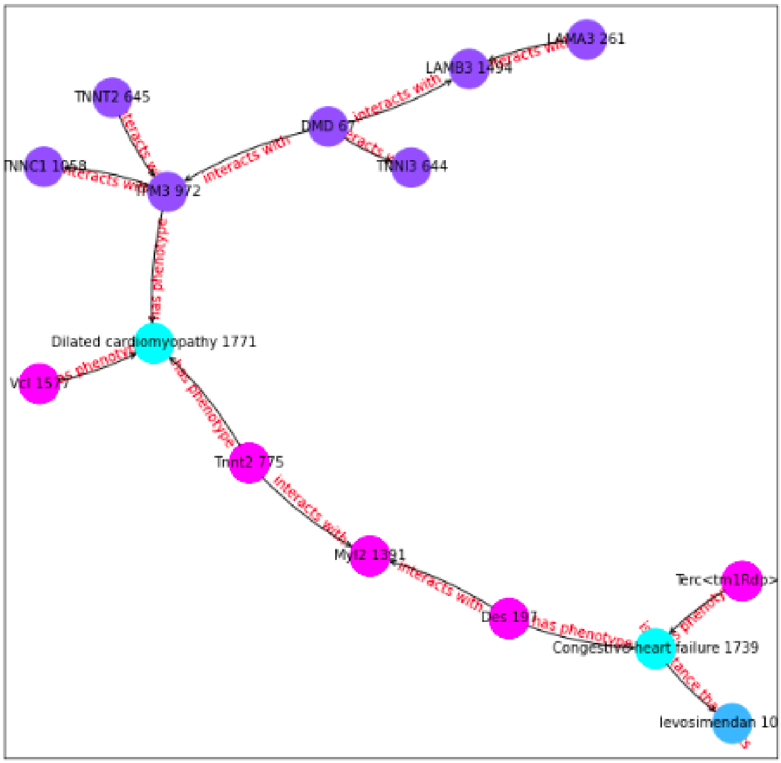


**Figure S8:** Explanation of drug candidate Levosimendan as possible treatment for Dilated cardiomyopathy. Classified as complete explanation.

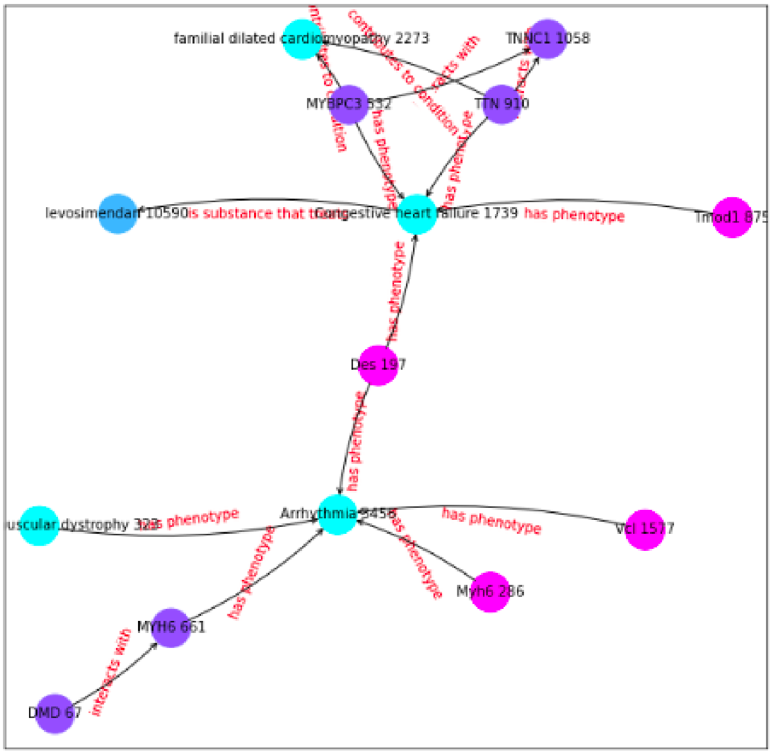


**Figure S9:** Explanation of drug candidate Levosimendan as possible treatment for Arrhythmia. Classified as complete explanation.

**Table S8**

Table showing the analysis of the explanation. The Good/Bad column shows the subjective evaluation. The Supporting Evidence Link shows if the Drug-Disease link contains supporting evidence. The Supporting Evidence Explanation shows if the explanation itself has supporting evidence.

| *Graph* | Drug | Disease | Good/Bad | Supporting Evidence Link | Supporting Evidence Explanation |
|---|---|---|---|---|---|
| *KG A* | Levosimendan | *Muscular Dystrophy* | Good | Yes | - |
| | Disopyramide | *Muscular Dystrophy* | Bad | Yes | - |
| | Entrectinib | *Muscular Dystrophy* | Good | No | - |
| | Entrectinib | *Respiratory Insufficiency* | Good | No | - |
| | Doxorubicin | *Respiratory Insufficiency* | Good | Contraindication | Unclear: `https://grantome.com/grant/NIH/R01-HL146443-01` |
| | Levosimendan | *Arrhythmia* | Bad | Yes | - |
| | Amiodarone | *Arrhythmia* | Good | Yes | - |
| | Isradipine | *Arrhythmia* | Good | Yes | - |
| | Levosimendan | *Dilated Cardiomyopathy* | Bad | Yes | - |
| | Aprindine | *Congestive Heart Failure* | Bad | Yes | - |
| | Nintedanib | *Congestive Heart Failure* | Good | No | - |
| | Levosimendan | *Progressive Muscle Weakness* | Good | Yes | `https://www.frontiersin.org/articles/10.3389/fphys.2021.786895/full` |
| | Entrectinib | *Cognitive Impairment* | Good | Contraindication | - |
| | Axitinib | *Cognitive Impairment* | Good | Yes | - |
| | Quercetin | *Cognitive Impairment* | Good | Yes | - |
| *KG B (Large)* | Methylprednisolone | *Muscular Dystrophy* | Good | Yes | - |
| | Methylprednisolone | *Respiratory Insufficiency* | Good | Yes | - |
| | Sorafenib | *Respiratory Insufficiency* | Good | Contraindication | Unclear: `https://www.ncbi.nlm.nih.gov/pmc/articles/PMC3961597/` |
| | Methylprednisolone | *Progressive Muscle Weakness* | Good | Contraindication | - |
| | Resveratrol | *Progressive Muscle Weakness* | Good | Yes | - |
| | Sorafenib | *Cognitive Impairment* | Good | Contraindication | - |

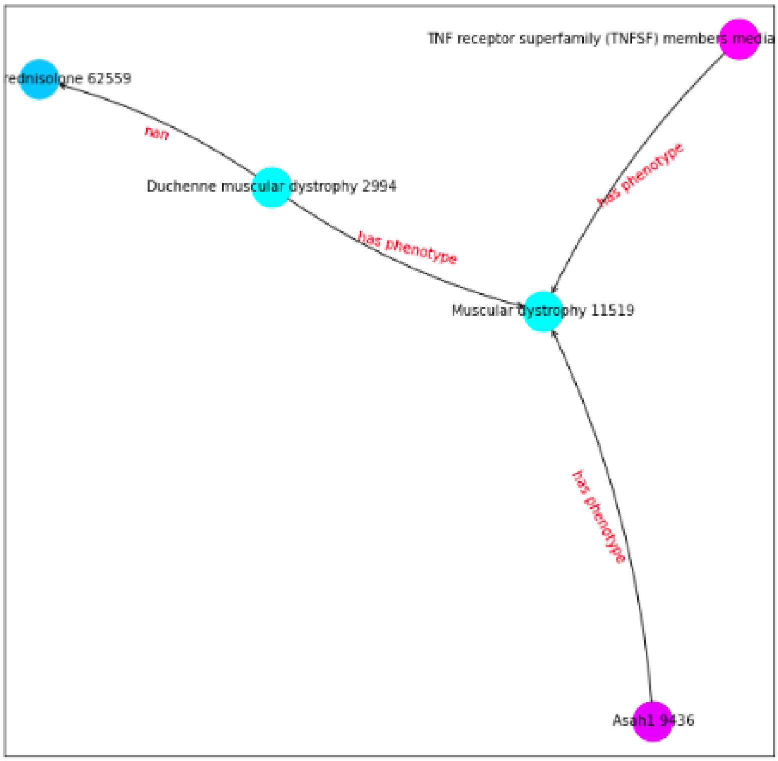


**Figure S10:** Explanation of drug candidate Methylprednisolone as possible treatment for Muscular dystrophy. Classified as complete explanation.

#### A.8. Drug Candidates on KG A

Table S9: Table showing the drug candidates with the highest scores for each symptom/phenotype obtained in Graph A. Any evidence that supports the prediction will be shown in the *Supporting Evidence* column. If the drug is contraindicated for the given symptom/phenotype it will also be shown in this column.

| Symptom | ID | Drug Candidate | Score | Supporting Evidence |
|---|---|---|---|---|
| Muscular dystrophy | HP:0003560 | Levosimendan | 0.849 | https://pubmed.ncbi.nlm.nih.gov/30796500/ |
| | | Disopyramide | 0.848 | https://pubmed.ncbi.nlm.nih.gov/7045292/ |
| | | Entrectinib | 0.845 | None |
| Respiratory insufficiency | HP:0002093 | Entrectinib | 0.954 | None |
| | | Axitinib | 0.925 | None |
| | | Doxorubicin | 0.915 | May produce respiratory dysfunction: https://grantome.com/grant/NIH/R01-HL146443-01 |
| Gowers sign | HP:0003391 | Entrectinib | 0.963 | None |
| | | Axitinib | 0.945 | None |
| | | Nintedanib | 0.932 | None |
| Global developmental delay | HP:0001263 | Entrectinib | 0.985 | Can produce developmental delay: https://www.ncbi.nlm.nih.gov/pmc/articles/PMC8341080/ |
| | | Axitinib | 0.974 | None |
| | | Nintedanib | 0.968 | None |
| Hyporeflexia | HP:0001265 | Entrectinib | 0.923 | None |
| | | Axitinib | 0.905 | None |
| | | Nintedanib | 0.872 | None |
| Proximal muscle weakness | HP:0003701 | Entrectinib | 0.961 | Can produce muscle weakness: https://www.drugs.com/sfx/entrectinib-side-effects.html |
| | | Axitinib | 0.944 | None |
| | | Nintedanib | 0.925 | https://pubmed.ncbi.nlm.nih.gov/29991677/ |
| Intellectual disability | HP:0001256 | Entrectinib | 0.947 | None |
| | | Axitinib | 0.921 | None |
| | | Doxorubicin | 0.884 | Can produce cognitive impairment: https://pubmed.ncbi.nlm.nih.gov/34055643 |
| Calf muscle pseudohypertrophy | HP:0003707 | Disopyramide | 0.813 | None |
| | | Entrectinib | 0.784 | None |
| | | Axitinib | 0.776 | None |
| Elevated serum creatine kinase | HP:0003236 | Entrectinib | 0.929 | Can increase more: https://www.oncolink.org/cancer-treatment/oncolink-rx/entrectinib-rozlytrek |
| | | Levosimendan | 0.920 | None |
| | | Disopyramide | 0.915 | None |
| Abnormal EKG | HP:0003115 | Levosimendan | 0.777 | https://pubmed.ncbi.nlm.nih.gov/20814559/ |
| | | Aprindine | 0.747 | https://pubmed.ncbi.nlm.nih.gov/10068848/ |
| | | Disopyramide | 0.713 | https://pubmed.ncbi.nlm.nih.gov/9141608/ |

| | | | | |
|---|---|---|---|---|
| Arrhythmia | HP:0011675 | Levosimendan | 0.890 | https://ccforum.biomedcentral.com/articles/10.1186/cc1595#:$\sim$:text=Effects%20of%20levosimendan%20on%20cardiac%20arrhythmia%20in%20patients%20with%20severe%20heart%20failure,-J%20Lilleberg%20%26&text=Levosimendan%20(LS)%20is%20a%20novel,oxygen%20consumption%2C%20and%20induces%20vasodilation. |
| | | Amiodarone | 0.792 | https://www.aafp.org/pubs/afp/issues/2003/1201/p2189.html#:$\sim$:text=Amiodarone%20is%20a%20potent%20antiarrhythmic,deaths%20in%20high%2Drisk%20patients. |
| | | Isradipine | 0.953 | https://pubmed.ncbi.nlm.nih.gov/8480504/ |
| Waddling gait | HP:0002515 | Entrectinib | 0.976 | None |
| | | Axitinib | 0.964 | None |
| | | Nintedanib | 0.947 | None |
| Dilated cardiomyopathy | HP:0001644 | Entrectinib | 0.967 | Can produce heart disease: https://www.drugs.com/cons/entrectinib.html |
| | | Levosimendan | 0.950 | https://pubmed.ncbi.nlm.nih.gov/25863426/#:$\sim$:text=Conclusions%3A%20Levosimendan%20seems%20to%20improve,support%20while%20awaiting%20heart%20transplantation. |
| | | Nintedanib | 0.933 | None |
| Flexion contracture | HP:0001371 | Entrectinib | 0.980 | None |
| | | Axitinib | 0.975 | None |
| | | Nintedanib | 0.958 | None |
| Specific learning disability | HP:0001328 | Entrectinib | 0.871 | None |
| | | Axitinib | 0.862 | None |
| | | Acepromazine | 0.830 | None |
| Skeletal muscle atrophy | HP:0003202 | Entrectinib | 0.962 | None |
| | | Axitinib | 0.946 | None |
| | | Nintedanib | 0.925 | https://pubmed.ncbi.nlm.nih.gov/29991677/ |
| Hypoventilation | HP:0002791 | Axitinib | 0.781 | None |
| | | Entrectinib | 0.769 | None |
| | | Mezlocillin | 0.759 | None |
| Calf muscle hypertrophy | HP:0008981 | Entrectinib | 0.978 | None |
| | | Axitinib | 0.977 | None |
| | | Disopyramide | 0.976 | None |
| Motor delay | HP:0001270 | Entrectinib | 0.991 | None |
| | | Sunitinib | 0.985 | None |
| | | Fedratinib | 0.978 | None |
| Generalized hypotonia | HP:0001290 | Entrectinib | 0.995 | None |
| | | Axitinib | 0.988 | None |
| | | Nintedanib | 0.983 | None |
| Cardiomyopathy | HP:0001638 | Levosimendan | 0.899 | https://www.ncbi.nlm.nih.gov/pmc/articles/PMC6588712/ |

| | | | | |
|---|---|---|---|---|
| | | Entrectinib | 0.848 | Can produce myocarditis: https://pubmed.ncbi.nlm.nih.gov/34315748/ |
| | | Carvedilol | 0.837 | https://www.ncbi.nlm.nih.gov/pmc/articles/PMC4055878/#:$\sim$:text=Pathways%20through%20which%20carvedilol%20exert,for%20beneficial%20effects%20in%20cardiomyopathy. |
| Hyperlordosis | HP:0003307 | Entrectinib | 0.970 | None |
| | | Axitinib | 0.959 | None |
| | | Disopyramide | 0.932 | None |
| Congestive heart failure | HP:0001635 | Entrectinib | 0.863 | Can produce heart failure: https://www.rozlytrek.com/ntrk/how-rozlytrek-may-help/possible-side-effects.html |
| | | Aprindine | 0.857 | https://pubmed.ncbi.nlm.nih.gov/6871919/ |
| | | Nintedanib | 0.835 | None |
| Delayed speech and language development | HP:0000750 | Entrectinib | 0.986 | None |
| | | Axitinib | 0.977 | None |
| | | Nintedanib | 0.969 | None |
| Scoliosis | HP:0002650 | Entrectinib | 0.994 | None |
| | | Axitinib | 0.989 | None |
| | | Nintedanib | 0.981 | None |
| Progressive muscle weakness | HP:0003323 | Levosimendan | 0.864 | https://www.frontiersin.org/articles/10.3389/fphys.2021.786895/full |
| | | Entrectinib | 0.985 | Can cause weakness: https://www.drugs.com/sfx/entrectinib-side-effects.html |
| | | Axitinib | 0.960 | Can cause weakness: https://www.mayoclinic.org/drugs-supplements/axitinib-oral-route/side-effects/drg-20075455?p=1#:$\sim$:text=This%20medicine%20may%20cause%20serious,trouble%20talking%2C%20or%20vision%20changes. |
| Cognitive impairment | HP:0100543 | Entrectinib | 0.952 | Can induce cognitive disorders: https://www.ncbi.nlm.nih.gov/pmc/articles/PMC8149347/#:$\sim$:text=Cognitive%20disorders%20included%20events%20reported,(0.2%25)%20%5B20%5D. |
| | | Axitinib | 0.931 | https://www.neuro-central.com/reversing-alzheimers-symptoms-in-mice-with-axitinib-treatment/ |
| | | Quercetin | 0.991 | https://www.ncbi.nlm.nih.gov/pmc/articles/PMC3736941/#:$\sim$:text=In%20vitro%20research%20also%20suggests,similar%20to%20that%20of%20caffeine. |

##### A.9. Drug Candidates on KG B

Table S10: Table showing the drug candidates with the highest scores for each symptom/phenotype obtained with Graph B. Any evidence that supports the prediction will be shown in the *Supporting Evidence* column. If the drug is contraindicated for the given symptom/phenotype it will also be shown in this column.

| Symptom | ID | Drug Candidate | Score | Reference |
|---|---|---|---|---|
| Muscular dystrophy | HP:0003560 | Methylprednisolone | 0.993 | https://pubmed.ncbi.nlm.nih.gov/17541998/ |
| | | Resveratrol | 0.963 | https://www.nature.com/articles/s41598-020-77197-6 |
| | | Tofisopam | 0.919 | https://extrapharmacy.ru/grandaxin-tofisopam-50mg-60tabs |
| Respiratory insufficiency | HP:0002093 | Methylprednisolone | 0.984 | https://jintensivecare.biomedcentral.com/articles/10.1186/s40560-018-0321-9 |
| | | Fedratinib | 0.981 | None |
| | | Sorafenib | 0.975 | Can cause pneumonia: https://www.ncbi.nlm.nih.gov/pmc/articles/PMC3961597/ |
| Gowers sign | HP:0003391 | Fedratinib | 0.994 | None |
| | | Bosutinib | 0.991 | None |
| | | Nintedanib | 0.990 | None |
| Global developmental delay | HP:0001263 | Fedratinib | 0.995 | None |
| | | Sorafenib | 0.994 | None |
| | | Bosutinib | 0.994 | None |
| Hyporeflexia | HP:0001265 | Fedratinib | 0.996 | None |
| | | Sunitinib | 0.994 | None |
| | | Bosutinib | 0.994 | None |
| Proximal muscle weakness | HP:0003701 | Fedratinib | 0.997 | Can produce muscle weakness: https://medlineplus.gov/druginfo/meds/a619058.html |
| | | Bosutinib | 0.995 | None |
| | | Methylprednisolone | 0.995 | Can produce weakness: https://erj.ersjournals.com/content/21/2/377.2#:$\sim$:text=Methylprednisolone%20is%20often%20given%20in,weakness%20following%20high%2Ddose%20steroids. |
| Intellectual disability | HP:0001256 | Fedratinib | 0.996 | None |
| | | Sorafenib | 0.995 | None |
| | | Bosutinib | 0.995 | None |
| Calf muscle pseudohypertrophy | HP:0003707 | Methylprednisolone | 0.970 | https://www.britannica.com/science/pseudohypertrophy |
| | | Ruxolitinib | 0.967 | https://www.sciencedirect.com/science/article/pii/S147148921630100X |
| | | Fedratinib | 0.948 | None |
| Elevated serum creatine kinase | HP:0003236 | Methylprednisolone | 0.994 | Can increase creatinine: https://www.ncbi.nlm.nih.gov/pmc/articles/PMC4275145/ |
| | | Fedratinib | 0.989 | Can increase more: https://jamanetwork.com/journals/jamaoncology/fullarticle/2330618 |
| | | Bosutinib | 0.982 | Can increase more: https://www.sciencedirect.com/science/article/pii/S2152265017305840 |

| | | | | |
|---|---|---|---|---|
| Abnormal EKG | HP:0003115 | Methylprednisolone | 0.982 | Can affect EKG: https://pubmed.ncbi.nlm.nih.gov/29668335/ |
| | | Patisiran | 0.879 | None |
| | | Silodosin | 0.878 | None |
| Arrhythmia | HP:0011675 | Methylprednisolone | 0.989 | Can produce arrhythmia: http://www.ijps.ir/article_2090.html#:$\sim$:text=Cardiac%20dysrhythmias%20have%20been%20reported,turn%2C%20may%20initiate%20cardiac%20dysrhythmias. |
| | | Fedratinib | 0.980 | None |
| | | Sorafenib | 0.979 | None |
| Waddling gait | HP:0002515 | Fedratinib | 0.991 | Can produce gait: https://www.accessdata.fda.gov/drugsatfda_docs/nda/2019/212327Orig1s000MultidisciplineR.pdf |
| | | Sorafenib | 0.990 | Can produce gait: https://www.ncbi.nlm.nih.gov/pmc/articles/PMC4094497/ |
| | | Midostaurin | 0.990 | None |
| Dilated cardiomyopathy | HP:0001644 | Methylprednisolone | 0.993 | https://pubmed.ncbi.nlm.nih.gov/25614863/ |
| | | Adefovir dipivoxil | 0.980 | None |
| | | Milrinone | 0.966 | https://pubmed.ncbi.nlm.nih.gov/10488574/#:$\sim$:text=Conclusion%3A%20Milrinone%20lactate%20is%20an,and%20IV%20of%20heart%20failure. |
| Flexion contracture | HP:0001371 | Fedratinib | 0.997 | None |
| | | Sorafenib | 0.996 | https://pubmed.ncbi.nlm.nih.gov/35274715/ |
| | | Bosutinib | 0.995 | None |
| Specific learning disability | HP:0001328 | Fedratinib | 0.984 | None |
| | | Sorafenib | 0.978 | None |
| | | Sunitinib | 0.977 | https://pubmed.ncbi.nlm.nih.gov/27046396/ |
| Skeletal muscle atrophy | HP:0003202 | Fedratinib | 0.995 | None |
| | | Ruxolitinib | 0.994 | None |
| | | Sunitinib | 0.993 | https://www.ncbi.nlm.nih.gov/pmc/articles/PMC4413636/ |
| Hypoventilation | HP:0002791 | Methylprednisolone | 0.990 | https://jintensivecare.biomedcentral.com/articles/10.1186/s40560-018-0321-9 |
| | | Resveratrol | 0.966 | None |
| | | Fedratinib | 0.993 | None |
| Calf muscle hypertrophy | HP:0008981 | Methylprednisolone | 0.978 | https://www.ncbi.nlm.nih.gov/pmc/articles/PMC2879072/ |
| | | Fedratinib | 0.977 | None |
| | | Resveratrol | 0.976 | https://journals.plos.org/plosone/article?id=10.1371/journal.pone.0083518 |
| Motor delay | HP:0001270 | Fedratinib | 0.995 | None |
| | | Sunitinib | 0.994 | https://www.ncbi.nlm.nih.gov/pmc/articles/PMC6586148/ |
| | | Vincristine | 0.993 | None |
| Generalized hypotonia | HP:0001290 | Fedratinib | 0.982 | None |

| | | | | |
|---|---|---|---|---|
| | | Sorafenib | 0.980 | None |
| | | Primidone | 0.980 | None |
| Cardiomyopathy | HP:0001638 | Methylprednisolone | 0.995 | https://pubmed.ncbi.nlm.nih.gov/7971647/ |
| | | Resveratrol | 0.974 | https://onlinelibrary.wiley.com/doi/full/10.1002/fsn3.92 |
| | | Adefovir dipivoxil | 0.971 | None |
| Hyperlordosis | HP:0003307 | Methylprednisolone | 0.986 | https://www.ncbi.nlm.nih.gov/pmc/articles/PMC4897302/ |
| | | Fedratinib | 0.982 | None |
| | | Sorafenib | 0.980 | None |
| Congestive heart failure | HP:0001635 | Methylprednisolone | 0.979 | https://www.sciencedirect.com/science/article/pii/S1071916414005843#:$\sim$:text=Methylprednisolone%20improved%20HF%20outcomes.,of%20patients%20from%20the%20study. |
| | | Daunorubicinol | 0.957 | Can produce cardiotoxicity: https://www.sciencedirect.com/topics/medicine-and-dentistry/daunorubicinol |
| | | Adefovir dipivoxil | 0.946 | None |
| Delayed speech and language development | HP:0000750 | Fedratinib | 0.994 | None |
| | | Midostaurin | 0.993 | None |
| | | Sunitinib | 0.993 | None |
| Scoliosis | HP:0002650 | Sorafenib | 0.995 | None |
| | | Fedratinib | 0.995 | None |
| | | Midostaurin | 0.994 | None |
| Progressive muscle weakness | HP:0003323 | Methylprednisolone | 0.999 | Can cause weakness: https://pubmed.ncbi.nlm.nih.gov/14629908/ |
| | | Resveratrol | 0.985 | https://pubmed.ncbi.nlm.nih.gov/33239684/ |
| | | Patisiran | 0.960 | None |
| Cognitive impairment | HP:0100543 | Sunitinib | 0.997 | None |
| | | Ruxolitinib | 0.997 | Can produce cognitive impairment: https://pubmed.ncbi.nlm.nih.gov/24661373/ |
| | | Bosutinib | 0.997 | https://pubmed.ncbi.nlm.nih.gov/34484904/ |

#### A.10. Drug Candidates AD

Table S11: Table showing the drug candidates with the highest scores for each symptom/phenotype obtained in the AD KG. Any evidence that supports the prediction will be shown in the *Supporting Evidence* column. If the drug is contraindicated for the given symptom/phenotype it will also be shown in this column.

| Symptom | Symptom ID | Candidate | Score | Evidence? |
|---|---|---|---|---|
| Personality changes | HP:0000751 | flortaucipir F 18 | 0.986 | https://www.sciencedirect.com/science/article/abs/pii/S0006322321015663 |
| | | fedratinib | 0.979 | May cause: https://medlineplus.gov/druginfo/meds/a619058.html |
| | | lansoprazole | 0.978 | None |
| Dysphagia | HP:0002015 | fedratinib | 0.998 | None |
| | | midostaurin | 0.998 | Causes no dysphagia ? https://www.ons.org/cjon/23/6/midostaurin-nursing-perspectives-managing-treatment-and-adverse-events-patients-flt3 |
| | | nintedanib | 0.997 | None |
| Alzheimer disease | HP:0002511 | Resveratrol | 0.983 | https://www.ncbi.nlm.nih.gov/pmc/articles/PMC5664214/ |
| | | pexidartinib | 0.980 | https://www.ncbi.nlm.nih.gov/pmc/articles/PMC8101105/ |
| | | memantine | 0.980 | https://pubmed.ncbi.nlm.nih.gov/16906789/ |
| Cerebral cortical atrophy | HP:0002120 | midostaurin | 0.998 | None |
| | | fedratinib | 0.998 | None |
| | | sunitinib | 0.998 | treats Brain Cancer: https://clinicaltrials.gov/ct2/show/NCT00923117 |
| Abnormality of extrapyramidal motor function | HP:0002071 | midostaurin | 0.991 | None |
| | | fedratinib | 0.990 | None |
| | | bosutinib | 0.989 | None |
| Dementia | HP:0000726 | midostaurin | 0.995 | https://www.sciencedirect.com/topics/chemistry/midostaurin |
| | | fedratinib | 0.995 | None |
| | | pazopanib | 0.994 | https://www.ncbi.nlm.nih.gov/pmc/articles/PMC5757517/ |
| Babinski sign | HP:0003487 | midostaurin | 0.998 | None |
| | | fedratinib | 0.997 | None |
| | | sunitinib | 0.997 | None |
| Lower limb hyperreflexia | HP:0002395 | flortaucipir F 18 | 0.964 | None |
| | | Donepezil | 0.956 | None |
| | | Clioquinol | 0.956 | None |
| Dysarthria | HP:0001260 | midostaurin | 0.999 | None |
| | | fedratinib | 0.999 | None |
| | | sunitinib | 0.998 | None |
| Memory impairment | HP:0002354 | flortaucipir F 18 | 0.982 | https://www.ncbi.nlm.nih.gov/pmc/articles/PMC8175307/ |
| | | pexidartinib | 0.977 | https://www.alzdiscovery.org/uploads/cognitive_vitality_media/Pexidartinib-Cognitive-Vitality-For-Researchers.pdf |
| | | sorafenib | 0.974 | None |
| Dystonia | HP:0001332 | fedratinib | 0.997 | None |

| | | | | |
|---|---|---|---|---|
| | | midostaurin | 0.997 | None |
| | | bosutinib | 0.995 | None |
| Optic ataxia | HP:0031868 | Clioquinol | 0.986 | None |
| | | Donepezil | 0.986 | None |
| | | Memantine | 0.970 | Optic nerve atrophy: https://pubmed.ncbi.nlm.nih.gov/26666888/ |
| Myoclonus | HP:0001336 | fedratinib | 0.996 | None |
| | | midostaurin | 0.996 | None |
| | | bosutinib | 0.996 | None |
| Apraxia | HP:0002186 | midostaurin | 0.989 | None |
| | | fedratinib | 0.988 | None |
| | | nintedanib | 0.988 | None |
| Seizure | HP:0001250 | fedratinib | 0.999 | Can cause: https://www.mskcc.org/cancer-care/patient-education/medications/fedratinib |
| | | midostaurin | 0.999 | None |
| | | bosutinib | 0.998 | Can cause: https://www.ema.europa.eu/en/documents/product-information/bosulif-epar-product-information_en.pdf |
| Gait disturbance | HP:0001288 | fedratinib | 0.986 | None |
| | | bosutinib | 0.977 | None |
| | | midostaurin | 0.973 | https://www.ncbi.nlm.nih.gov/pmc/articles/PMC8301989/ |
| Neurofibrillary tangles | 0.973 | flortaucipir F 18 | 0.974 | https://pubchem.ncbi.nlm.nih.gov/compound/70957463 |
| | | cycloserine | 0.962 | https://pubmed.ncbi.nlm.nih.gov/36159454/ |
| | | lansoprazole | 0.961 | https://pubmed.ncbi.nlm.nih.gov/24900410/ |
| Spastic tetraparesis | HP:0001285 | duloxetine | 0.959 | None |
| | | flortaucipir F 18 | 0.952 | None |
| | | metformin | 0.951 | None |
| Agnosia | HP:0010524 | Donepezil | 0.980 | https://www.ncbi.nlm.nih.gov/pmc/articles/PMC3504981/ |
| | | Clioquinol | 0.980 | None |
| | | Memantine | 0.967 | https://pubmed.ncbi.nlm.nih.gov/19898670/ |

#### A.11. Drug Candidates ALS

Table S12: Table showing the drug candidates with the highest scores for each symptom/phenotype obtained in the ALS KG. Any evidence that supports the prediction will be shown in the *Supporting Evidence* column. If the drug is contraindicated for the given symptom/phenotype it will also be shown in this column.

| Symptom | Symptom ID | Candidate | Score | Evidence? |
|---|---|---|---|---|
| Sleep apnea | HP:0010535 | Riluzole | 0.808 | https://pubmed.ncbi.nlm.nih.gov/11732759/ |
| | | Gabapentin | 0.767 | Can cause: https://pubmed.ncbi.nlm.nih.gov/28116804/ |
| | | Vitamin E | 0.756 | https://pubmed.ncbi.nlm.nih.gov/23389837/ |
| Degeneration of anterior horn cells | HP:0002398 | Riluzole | 0.821 | Spinal muscular atrophy: https://pubmed.ncbi.nlm.nih.gov/14623733/ |
| | | tacrolimus | 0.785 | Not significant: https://www.nature.com/articles/sc2015172 |
| | | brilliant Blue G | 0.768 | Can help ELA: https://peerj.com/articles/3064/ |
| Dysarthria | HP:0001260 | hexachlorophene | 0.976 | None |
| | | dabrafenib | 0.972 | None |
| | | dichlorophen | 0.954 | None |
| Skeletal muscle atrophy | HP:0003202 | hexachlorophene | 0.953 | None |
| | | dabrafenib | 0.935 | Can cause: https://pubmed.ncbi.nlm.nih.gov/32898388/ |
| | | quercetin | 0.907 | https://pubmed.ncbi.nlm.nih.gov/25614714/#:$\sim$:text=Together%2C%20these%20findings%20suggest%20that,induced%20muscle%20inflammation%20and%20sarcopenia. |
| Muscle weakness | HP:0001324 | hexachlorophene | 0.951 | None |
| | | dabrafenib | 0.944 | None |
| | | quercetin | 0.989 | https://www.ncbi.nlm.nih.gov/pmc/articles/PMC6356612/#:$\sim$:text=Taken%20together%2C%20the%20findings%20from,sarcolemmal%20action%20potential%20propagation%20impairment. |
| Muscle spasm | HP:0003394 | hexachlorophene | 0.975 | None |
| | | dabrafenib | 0.958 | Can cause: https://www.macmillan.org.uk/cancer-information-and-support/treatments-and-drugs/dabrafenib-and-trametinib |
| | | dichlorophen | 0.945 | None |
| Amyotrophic lateral sclerosis | HP:0007354 | hexachlorophene | 0.911 | https://pubmed.ncbi.nlm.nih.gov/25987361/ |
| | | oleic acid | 0.884 | https://pubmed.ncbi.nlm.nih.gov/29760648/ |
| | | dabrafenib | 0.881 | None |

| | | | | |
|---|---|---|---|---|
| Dysphagia | HP:0002015 | hexachlorophene | 0.987 | None |
| | | dabrafenib | 0.980 | None |
| | | dichlorophen | 0.968 | None |
| Fasciculations | HP:0002380 | hexachlorophene | 0.920 | None |
| | | oleic acid | 0.891 | Can increase: https://www.sciencedirect.com/science/article/pii/S0006899314005861?via%3Dihub |
| | | dabrafenib | 0.998 | None |
| Degeneration of the lateral corticospinal tracts | HP:0002314 | hexachlorophene | 0.829 | None |
| | | dabrafenib | 0.787 | None |
| | | celecoxib | 0.787 | None |
| Pseudobulbar paralysis | HP:0007024 | Riluzole | 0.788 | https://www.nejm.org/doi/full/10.1056/NEJM199403033300901 |
| | | Gabapentin | 0.727 | None |
| | | celecoxib | 0.720 | None |
| Hyperreflexia | HP:0001347 | hexachlorophene | 0.983 | Can cause: https://pubchem.ncbi.nlm.nih.gov/compound/Hexachlorophene#section=Human-Toxicity-Excerpts |
| | | dabrafenib | 0.977 | None |
| | | dichlorophen | 0.963 | None |
| Spasticity | HP:0001257 | hexachlorophene | 0.989 | None |
| | | dabrafenib | 0.986 | None |
| | | sotorasib | 0.970 | None |
